# Speed tuning in the macaque posterior parietal cortex

**DOI:** 10.1101/204966

**Authors:** Eric Avila, Kaushik J Lakshminarasimhan, Gregory C DeAngelis, Dora E Angelaki

## Abstract

Neurons in the macaque posterior parietal cortex are known to encode the *direction* of self-motion. But do they also encode one’s *speed?* To test this, we performed neural recordings from area 7a while monkeys were passively translated or rotated at various speeds. Visual stimuli were delivered as optic flow fields and vestibular stimuli were generated by a motion platform. Under both conditions, the responses of a fraction of neurons scaled linearly with self-motion speed, and speed-selective neurons were not localized to specific layers or columns. We analyzed ensembles of simultaneously recorded neurons and found that the precision of speed representation was sufficient to support path integration over modest distances. Our findings describe a multisensory neural code for linear and angular self-motion speed in the posterior parietal cortex of the macaque brain, and suggest a potential role for this representation.

## INTRODUCTION

Efficient navigation demands the use of static, external sensory cues as well as dynamic cues generated by self-motion. Animals rely mostly on external landmarks to navigate in familiar environments. However when landmarks are unavailable, they are capable of using self-motion signals to navigate by path integration (see Etienne & Jeffery, 2004 for review). Path integration entails maintaining an estimate of one’s position by integrating linear and angular velocities. Consistent with this, the amplitude and phase of response fields of spatially tuned neurons in entorhinal cortex and hippocampus are modulated by animals’ velocity (Sargolini et al., 2006; Lu and Bilkey, 2010; Yoon et al., 2013;Cowen and Nitz, 2014). Yet the precise source of velocity inputs to the navigation circuit in primates is unclear.

A candidate cortical area that might convey velocity information to navigation circuits is the posterior parietal cortex (PPC). PPC plays a major role in directing spatial behavior (Cohen and Andersen, 2002; Save and Poucet, 2008; Nitz, 2009), and rats with PPC lesions exhibit impaired spatial navigation (Kolb et al., 1994; Save and Poucet, 2000) and inaccurate path integration (Parron and Save, 2004; Save, et al., 2001). Furthermore, a growing body of evidence suggests that a large fraction of neurons in rat PPC is modulated by the animals’ locomotor state (McNaughton et al., 1994; Nitz, 2006) including movement speed (Whitlock et al., 2012; Wilber et al., 2017). In contrast, the tuning of PPC neurons to allocentric variables (e.g., route information or landmarks) is weak at best (Chen et al., 1994a; Chen et al., 1994b).

The above findings are yet to be fully corroborated in primates. The macaque PPC has a relatively more complex parcellation and anatomical connectivity than rodent PPC (e.g. Kaas et al., 2013). Although neuronal responses in parietal areas such as MSTd and VIP to multisensory self-motion stimuli are well documented (Bremmer et al., 2002(a,b); Britten and van Wezel, 1998; Britten and Van Wezel, 2002; Chen et al., 2011; Gu et al., 2006; Maciokas and Britten, 2010; Zhang and Britten, 2010), these are not the areas with the strongest connectivity to navigation circuits. Anatomical studies suggest that the posterior aspect of the inferior parietal gyrus, between the intraparietal and superior temporal sulci (area 7a), constitutes the main region of PPC with both direct and indirect connections to the hippocampal formation (Andersen et al., 1990; Ding et al., 2000; Cavada and Goldman-Rakic, 1989; Kobayashi and Amaral, 2000, 2003; Kondo et al., 2005; Morris et al., 1999; Pandya and Seltzer, 1982; Rockland and Van Hoesen, 1999; Vogt and Pandya, 1987). The indirect pathway via the retrosplenial cortex has been proposed to play a critical role in communicating navigationally relevant information (Kravitz et al., 2011; Burgess et al., 1999). Several studies appear to support this view. First, lesions to area 7a in cynomolgus monkeys impaired navigation of whole-body translation through a maze (Traverse and Latto, 1986; Barrow and Latto, 1996). Second, motion-sensitive receptive fields of area 7a neurons often exhibit a radial arrangement of preferred directions within their receptive field (Motter and Mountcastle, 1981; Steinmetz et al., 1987), such that they are maximally activated by the expanding or contracting patterns of optic flow that are typically experienced when the observer moves forward or backward through the world (Siegel and Read, 1997; Merchant et al., 2001, 2003; Raffi and Siegel, 2007). Finally, the response gain of these neurons is also modulated by eye/head position (‘gain fields’, Andersen et al., 1985), suggesting that they may play a role in transforming sensory inputs into a navigationally useful format such as a body-centered representation (Zipser and Andersen, 1988; Pouget and Sejnowski, 1997). Thus, anatomical and physiological evidence suggests that area 7a plays a role in bridging motion perception and navigation. Yet, no study has explicitly tested whether 7a neurons represent linear and angular velocities necessary for self-motion-based navigation. Moreover, while responsiveness to optic flow has been described (Phinney and Siegel, 2000; Raffi and Siegel, 2007; Read and Siegel, 1997), responses of 7a neurons to vestibular stimulation have never been directly recorded.

We recorded neuronal spikes and local-field potentials in area 7a during forward translation as well as yaw rotation paradigms to probe the neural representation of linear and angular velocity. We systematically varied speed using both visually simulated (optic flow) and passive vestibular (motion platform) self-motion and found two key results. First, contrary to the general belief that responses in area 7a are predominantly visual, we found evidence for a vestibular dominance in self-motion processing. Second, similar to rodent PPC, neural response increased with the magnitude of linear and angular velocity. Our results suggest that area 7a carries a multisensory code for self-motion velocity and may play a role in path integration.

## RESULTS

### Response to linear translation

We recorded local field potentials (LFP) from 704 channels across 44 recording sessions and isolated 340 neurons from area 7a of three macaque monkeys while they passively fixated and experienced real and/or visually-simulated forward motion at different speeds (**Methods, Fig. 1A, Supplementary Fig. S1**). All recordings were carried out extracellularly using laminar electrodes with 16 contact sites, spaced 100μm apart, allowing us to sample neural activity spanning 1.5 mm across the depth of the cortex (**Fig. 1B**).

We first examined stimulus-induced fluctuations in the LFP. The shape of the LFP waveforms was highly correlated across channels (mean pairwise Pearson’s correlation *r=0.84±0.1*), as was the timing and amplitude of their peak fluctuations (**Supplementary Fig. S2**). Therefore, we averaged LFP traces from different electrodes and different speeds together before analyzing stimulus-evoked responses for each condition. We found that evoked potentials were highly stereotyped across recording sessions (**Fig. 1C**). The onset of motion induced a large change in LFP, whose amplitude and latency both depended on stimulus condition. Across all recordings, vestibular LFP responses had significantly greater peak amplitudes, defined as the largest deflection away from baseline during the motion period, as well as lower peak latencies than responses evoked by visual motion (mean ± standard-error, vestibular: 121±3 μV at 214±1 ms; visual: 42±2 μV at 238±3 ms; *p*=4x10^−94^, *t*-test for difference in amplitudes; *p*=2x10^−22^, *t*-test for difference in latencies; *n*=704 sites). Furthermore, the amplitude of evoked potentials elicited by the combination of vestibular and visual stimuli (combined: 122±4 μV at 211±1 ms; *n*=704 sites) was much more strongly correlated with the vestibular response (Pearson’s correlation, *r* = 0.89, *p*=2x10^−245^ than the visual response (*r* = 0.16, *p*=9x10^−6^). These findings show that area 7a is responsive to forward motion, and is likely to be more sensitive to vestibular than visual motion cues.

**Figure 1.**
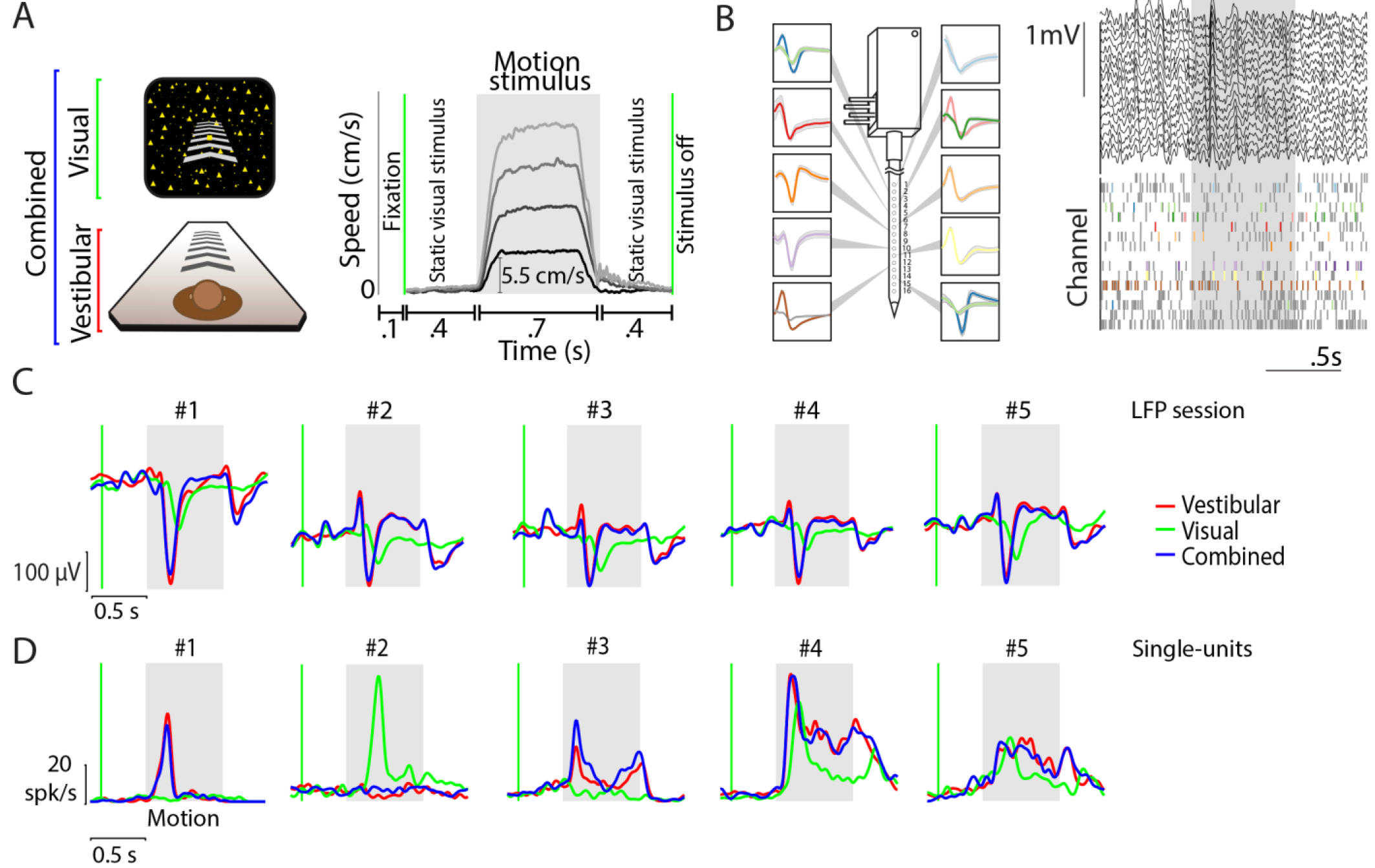
Example responses to linear translation. **(A)** *Left-stimulus conditions*: Monkeys were seated on a motion platform in front of a projector screen, and experienced simulated forward translation via optic flow (visual), real forward translation delivered using the platform (vestibular) or a congruent combination of the two (combined) while fixating on a central target. *Right-trial structure*: Time course of translation velocity reconstructed from accelerometer recordings during four representative trials under the vestibular condition. The desired temporal profile of velocity was trapezoidal for both vestibular and visual motion, and one of four possible movement speeds (5.5, 11, 16.5, 22 cm/s) was presented on each trial. During trials with visual motion, the motion period was flanked by the presentation of a static cloud of dots for 400 ms. No visual stimulus, except for the fixation target, was present during these periods under the vestibular condition. **(B)** Left: Recordings were carried out using a linear electrode array (*U-Probe*) with 16 contact sites (waveforms illustrate recorded spikes from an example session; shaded regions show ± 1 standard deviation). Right: Local field potentials (LFP, top) and spikes (bottom) from multi-units (grey) and isolated single-units (colored) during one trial. **(C)** Stimulus-evoked LFP (averaged across all channels of the array and all stimulus speeds) during vestibular (red), visual (green), and combined (blue) conditions of five example recording sessions carried out at different locations. Grey shaded areas correspond to periods of movement. **(D)** Response time-course of five example neurons, averaged across all speeds, chosen to highlight the diversity in temporal dynamics within the population. Green vertical line shows start of the static cloud of dots for the visual and combined conditions. Grey shaded region denotes the motion period. After motion offset, the monkey maintained fixation for another 400 ms (and the static cloud of dots was maintained for visual and combined conditions).

In contrast to the LFP, the time-course of responses of individual neurons was quite diverse even among neurons recorded simultaneously at different depths along the same penetration (**Supplementary Fig. S3**). **Figure 1D** shows some example temporal response patterns. Whereas some neurons responded with a transient change in activity following onset and/or offset of the motion stimulus, a large number of neurons exhibited a change that persisted throughout the period of motion. There was also a notable diversity in the polarity of response: although the vast majority of responsive neurons exhibited an increase in firing rate following motion onset (‘excitatory’), a small but significant fraction of neurons was suppressed by motion (‘suppressive’, Methods). Despite this heterogeneity, some key aspects of neuronal response were consistent with LFP responses. Specifically, a larger fraction of neurons was significantly responsive to vestibular (~40%) than visual motion (~27%, Table 1A). Moreover, the time-course of neuronal activity during combined visual-vestibular motion was much more strongly correlated with that of the vestibular response (mean pairwise Pearson’s correlation, *r* = 0.82 ± 0.25) than the visual response (*r* = 0.57 ± 0.2), suggesting that neuronal response dynamics to multisensory inputs are more strongly dictated by vestibular rather than visual inputs. We asked whether there was true multisensory convergence of visual and vestibular self-motion signals at the neuronal level, that is, whether vestibular-responsive neurons are likely to also respond to visual motion and vice-versa. Although the proportion of neurons responding to the combined stimulus was relatively large (~38%), only a small subset of those neurons responded to both individual conditions when presented separately (~17%). In fact, given the proportion of vestibular and visual neurons in our population, the number of bi-sensory neurons (*n*=57) was only marginally greater than expected by chance (99% confidence interval for the number of bi-sensory neurons: *n*=[25,51]). Therefore, the observed fraction of bi-sensory neurons can largely be explained by independent, rather than cooperative, convergence of vestibular and visual inputs, providing only weak evidence for multisensory convergence of motion information at the neuronal level.

**Table 1.** Statistics of neuronal and multi-unit response to linear speed. **(A)** Single-units were classified into two groups based on whether at least one of the linear speeds elicited response that was significantly higher (excitatory) or lower (suppressive) response than their baseline response (fixation-only trials) (*t*-test corrected for multiple comparisons; significance-level *p* = 0.05). A subset of responsive neurons were identified as ‘speed tuned’ to linear motion if their responses were significantly different across speeds (one-way ANOVA; *p* = 0.05). Values between brackets correspond to percentages of the total number of recorded neurons. **(B)** Corresponding statistics for multi-unit responses to linear translation.

| A Single-unit |  |  |  | B Multi-unit |  |  |  |
| --- | --- | --- | --- | --- | --- | --- | --- |
| Neuron summary | Excitatory | Suppressive | Total | Neuron summary | Excitatory | Suppressive | Total |
| Vestibular | 118(34.7) | 15(4.4) | 133(39.1) | Vestibular | 530(75.2) | 23(3.2) | 553(78.5) |
| Visual | 77(22.6) | 15(4.4) | 92(27) | Visual | 235(33.3) | 66(9.3) | 301(42.7) |
| Combined | 124(36.4) | 6(1.7) | 130(38.2) | Combined | 520(73.8) | 13(1.8) | 533(75.7) |
| Vestibular and visual | 49(14.4) | 8(2.3) | 57(16.7) | Vestibular and visual | 198(28.1) | 3(0.4) | 201(28.5) |
| Speed Tuned | Excitatory |  |  | Speed Tuned | Excitatory |  |  |
| Vestibular | 52(15.2) |  |  | Vestibular | 215(30.5) |  |  |
| Visual | 25(7.3) |  |  | Visual | 57(8.1) |  |  |
| Combined | 51(15) |  |  | Combined | 240(34) |  |  |
| Vestibular and visual | 11(3.2) |  |  | Vestibular and visual | 43(6.1) |  |  |

**Figure 2.**
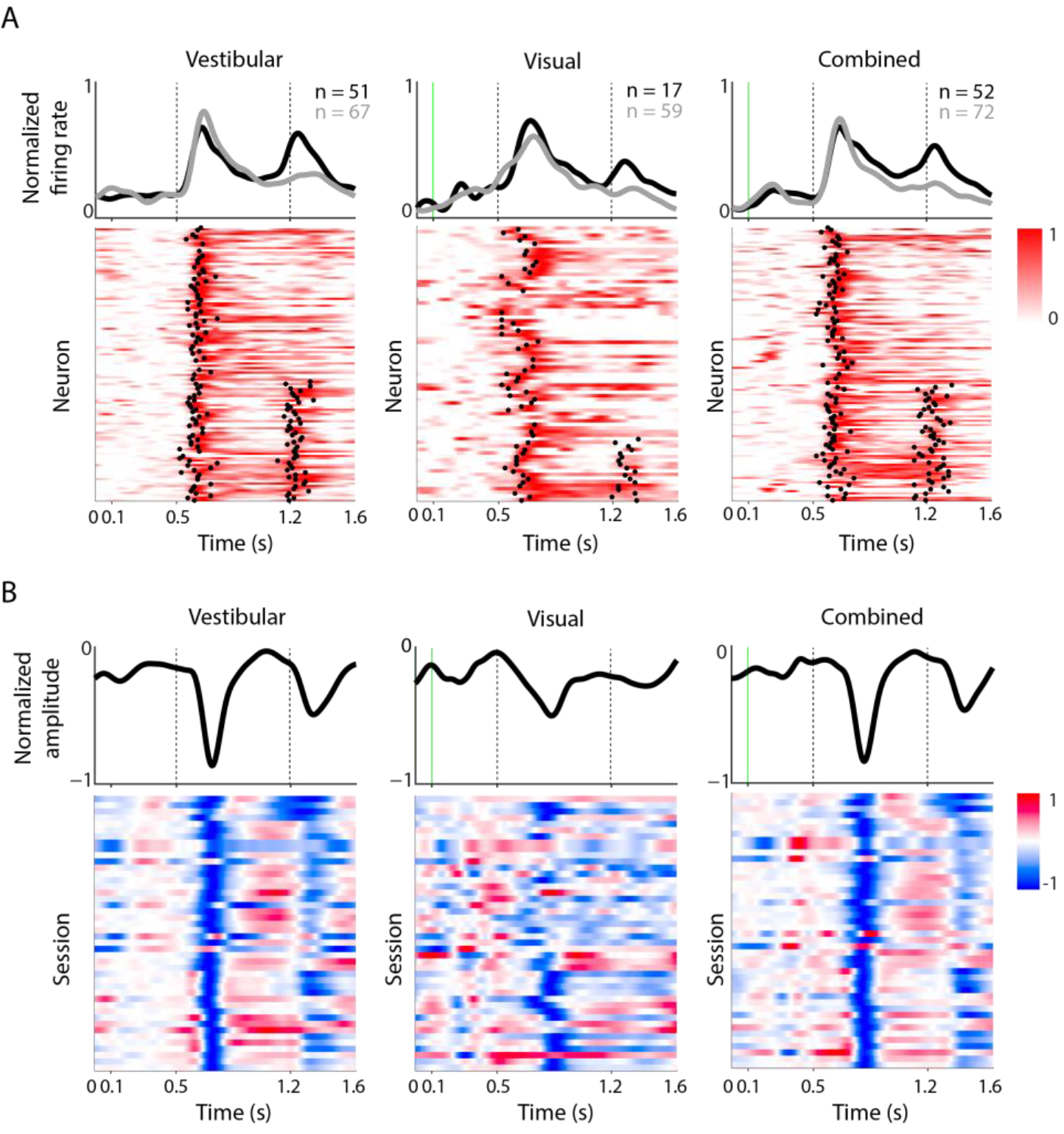
Temporal pattern of response. **(A)** The rows in each panel correspond to normalized firing rates of all neurons that increased their firing (‘excitatory’) to vestibular (*left*), visual (*middle*), and combined (*right*) motion. To normalize the responses, we subtracted the mean pre-stimulus activity of each neuron from its response, and then divided by its peak response. We classified the temporal responses as unimodal or bimodal depending on the number of time-points at which a sudden increase in response was detected (black dots-Methods). Both unimodal (following motion onset) and bimodal (following motion onset/offset) profiles are frequently encountered in response to vestibular and combined motion. In contrast, responses are mostly unimodal when only visual motion is present. Responses of neurons with unimodal (gray) and bimodal (black) profiles are averaged separately and shown on top of the respective panels. Vertical dotted lines show motion onset and offset. **(B)** Response profiles of LFPs (averaged across all channels) for all recorded sessions (*n*=44), normalized in the same way as the single-unit responses. Note that LFPs were strongly negative at most sites.

Figure 2A shows response dynamics of the set of all responsive neurons that exhibited a motion-induced increase (‘excitatory’) in firing rate within 400 ms of motion onset. An analysis of these dynamics (Methods) revealed two distinct populations. During vestibular motion, the response profile of nearly half of the neurons (~43%) was bimodal, exhibiting an increase after both the onset and offset of motion (**Fig. 2A** - left). This fraction was not altered by the addition of visual motion cues (42%, **Fig. 2A** - right). In contrast, responses to purely visual motion were mostly unimodal (Fig. 2A - middle). Vestibular responses had significantly shorter latencies than visual responses (vestibular: 128±5 ms, visual: 140±6 ms; *t*-test, *p*=0.019). The latency of responses to the combined stimulus was more strongly correlated with vestibular (r=0.37, *p*= 4.4x10^−4^) than visual (r=0.007, *p*=0.96) latencies. Moreover, the peak firing rates of the subset of neurons responsive to combined stimuli were more strongly modulated by vestibular than visual cues (**Supplementary fig. S4**). These results parallel those of the LFPs and reinforce the view that the neural representation of linear motion in area 7a is strongly influenced by vestibular inputs.

Very similar results were found for multi-unit activity (MUA). Of 704 recordings sites, MUA at 553 (~78%) sites were significantly responsive to vestibular input, 301 (~43%) sites were responsive to visual input, and 533(~76%) sites were responsive to the combined stimulus (**Table 1B**). Similar to single-units, MUA exhibited diverse temporal dynamics (**Supplementary Fig. S5**) with nearly half of recordings showing an increase in firing after both the onset and offset of vestibular motion (**Supplementary Fig. S6**). Moreover, multi-unit responses were correlated strongly with those of single-units recorded from the same electrode site (**Supplementary Fig. S7A**). Consequently, multi-units also exhibited stronger modulation to vestibular than visual inputs (**Supplementary Fig. S7B**).

**Figure 3.**
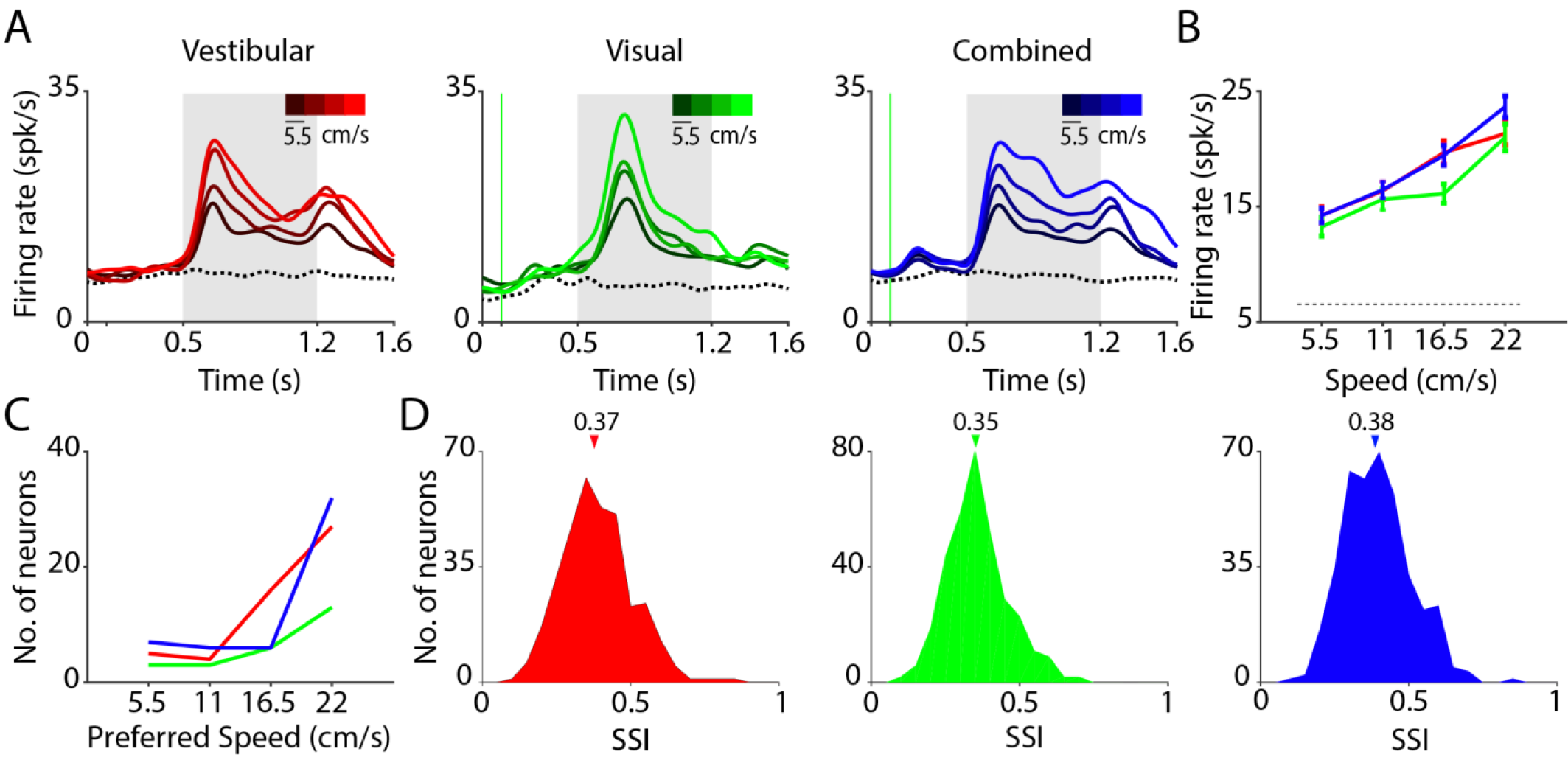
Translation speed modulates neuronal responses in 7a. **(A)** Time course of responses averaged across neurons that were significantly tuned to linear speed in the vestibular (*left, n=52*), visual (*middle, n=25*), and combined (*right, n=51*) conditions. Trials are grouped and color-coded according to motion speed (brighter hues correspond to greater speeds). Dotted trace denotes response to the baseline condition (fixation only trials). Grey shaded areas correspond to periods of movement. **(B)** Average tuning curves of single-units in (A) to translation speed under the three conditions (red-vestibular, green-visual, blue-combined). **(C)** Distribution of preferred speed (the speed that elicited the greatest firing rate during the flat phase of the trapezoidal motion profile) for neurons that exhibited excitatory responses to motion. **(D)** Distribution of the speed selectivity index (*SSI*) of all neurons in the three conditions. Arrows on top indicate the medians of the distributions.

We tested whether neurons responsive to translational motion also encoded translation speed (**Methods**). Since the vast majority of the responsive neurons were excited (rather than suppressed) by motion (**Table 1A**, top), we restricted our analysis to this group of neurons. We found that many of these neurons were tuned to speed (Vestibular: *n*=52/118; Visual: *n*=25/77; Combined: *n*=51/124; ANOVA, *p*<0.05; **Table 1A**, bottom). To assess the linear trend in neuronal tuning for speed, we pooled the responses of all tuned neurons separately to each movement speed, and estimated the correlation with translation speed. Across the population of tuned neurons, the responses were positively correlated with speed under all stimulus conditions (vestibular: *r*=0.21, *p*=2x10^−3^; visual: *r*=0.27, *p*=5x10^−3^; combined: *r*=0.30, *p* =1.6x10^−5^), as readily seen from the population average of response time-courses and tuning functions (**Fig. 3A, B**). Consequently, the distribution of preferred speed, defined as the speed that elicited the largest deflection from baseline, was skewed towards higher speeds (**Fig. 3C**). Analysis of multiunit responses yielded very similar results (**Supplementary Fig. S8**).

To quantify the sensitivity of each neuron to speed, we computed a speed selectivity index (SSI), based on its response to different speeds (**Methods**). Comparing the distribution of SSIs across all neurons (**Fig. 3D**), we found that the median SSI value for the vestibular condition (0.37±0.11) was slightly but significantly greater than that for the visual condition (0.35±0.10, *p*=1.6x10^−5^, paired *t*-test), suggesting that speed selectivity is greater, overall, under the vestibular condition (**Supplementary Fig. S9**). Moreover, the median SSI under the combined condition was significantly greater than that of the visual (paired *t*-test, *p*=1.4x10^−7^) but not the vestibular (*t*-test, *p*=0.28) condition, suggesting that speed tuning in the combined condition is more likely to be driven by vestibular inputs.

Finally, we analyzed the spike waveforms to identify neuronal subtypes (**Methods, Supplementary Fig. S10**) and found a bimodal distribution of spike widths (Hartigan’s dip-test, *p* = 0.036, Hartigan and Hartigan, 1985). On average, the response dynamics and speed selectivity of the putative broad- and narrow-spiking neurons were similar, suggesting that putative pyramidal and interneurons may share the same code.

### Response to angular rotation

If area 7a contains a code for self-motion necessary for path integration, neurons should also signal angular velocity. We recorded from 268 isolated neurons (across 36 recording sessions) using a protocol in which the motion stimulus was composed of a pure yaw rotation at one of several angular velocities (**Methods, Fig. 4A**), including both clockwise (CW) and counter-clockwise (CCW) rotation. These stimuli evoked changes in the LFP that were very similar to those triggered by linear translation. The onset of inertial as well as visually-simulated rotation both resulted in reliable evoked potentials that were highly correlated across recording sites at different depths (*r=0.84±0.1*, **Fig. 4B**, top and **Supplementary Fig. S11**), and across different recording sessions (**Fig. 4C**). Vestibular rotation produced larger and faster responses in the LFP than visual rotation (mean ± standard-error, vestibular: 93+3 μV at 218±1 ms; visual: 37+2 μV at 253±3 ms; *p*=1.4x10^−50^, *t*-test for difference in amplitude; *p*=2.5x10^−14^, *t*-test for difference in latency; **Fig. 4C**). Combined visual-vestibular rotation was not included in this protocol due to technical limitations (**Methods**).

Single- and multi-unit responses were also similar to those observed in response to translational motion in that there was a diverse pattern of temporal responses (**Fig. 4D, Supplementary Figs. S12, S13**). Both vestibular and visual rotational motion elicited responses in a large fraction of neurons (**Table 2A**; ~31% for vestibular, ~20% for visual motion). Nevertheless, consistent with the findings from translational motion, the number of multisensory neurons (*n*=29) barely exceeded that expected by chance (*99^th^* percentile CI: [10,27]), again suggesting weak cue-integration at the single neuron level. The average latency of neuronal responses to vestibular rotation shows a slight tendency toward faster responses, as compared with visual rotation (vestibular: 124±5 ms; visual: 143±10 ms; *p*=0.07, *t*-test). These results corroborate findings from the linear translation protocol, and suggest that area 7a is more sensitive to vestibular cues both for translational and rotational motion.

For each stimulus condition, we found that a significant proportion of neurons that were excited by rotational motion showed significant tuning (Vestibular: *n*=26/75; Visual: *n*=27/51, ANOVA, *p*<0.05) for the speed of rotation (**Table 2A, Fig. 5A, B**). We tested the direction selectivity of individual neurons by comparing responses from trials with clockwise rotation against responses from trials with counterclockwise rotation. We found that only a small fraction of neurons was significantly tuned to direction of rotation (Vestibular: *n*=4/75; Visual: *n*=9/51; *p*<0.05, *t*-test). Consequently, neuronal responses to the fastest speed of CW and CCW rotation were strongly correlated across the population (Vestibular: *r*=0.87, *p*=4.57x10^−85^; Visual: *r*=0.77, *p*=2.65x10^−54^, **Fig. 5C**). This indicates that 7a neurons largely signal the magnitude but not the direction of rotational velocity. We estimated the correlation between neuronal responses and angular speed by pooling trial-averaged responses of all neurons to each rotation speed separately (**Methods**). Since neural responses were largely unaffected by rotation direction, we combined responses from CW and CCW directions before trial averaging. We observed a significant positive correlation between neuronal activity and rotation speed in the vestibular condition (Pearson’s correlation *r*=0.18, *p*=0.004). A similar tendency was present for the visual condition, although the correlation did not quite reach statistical significance (*r*=0.07, *p*=0.06).

**Figure 4.**
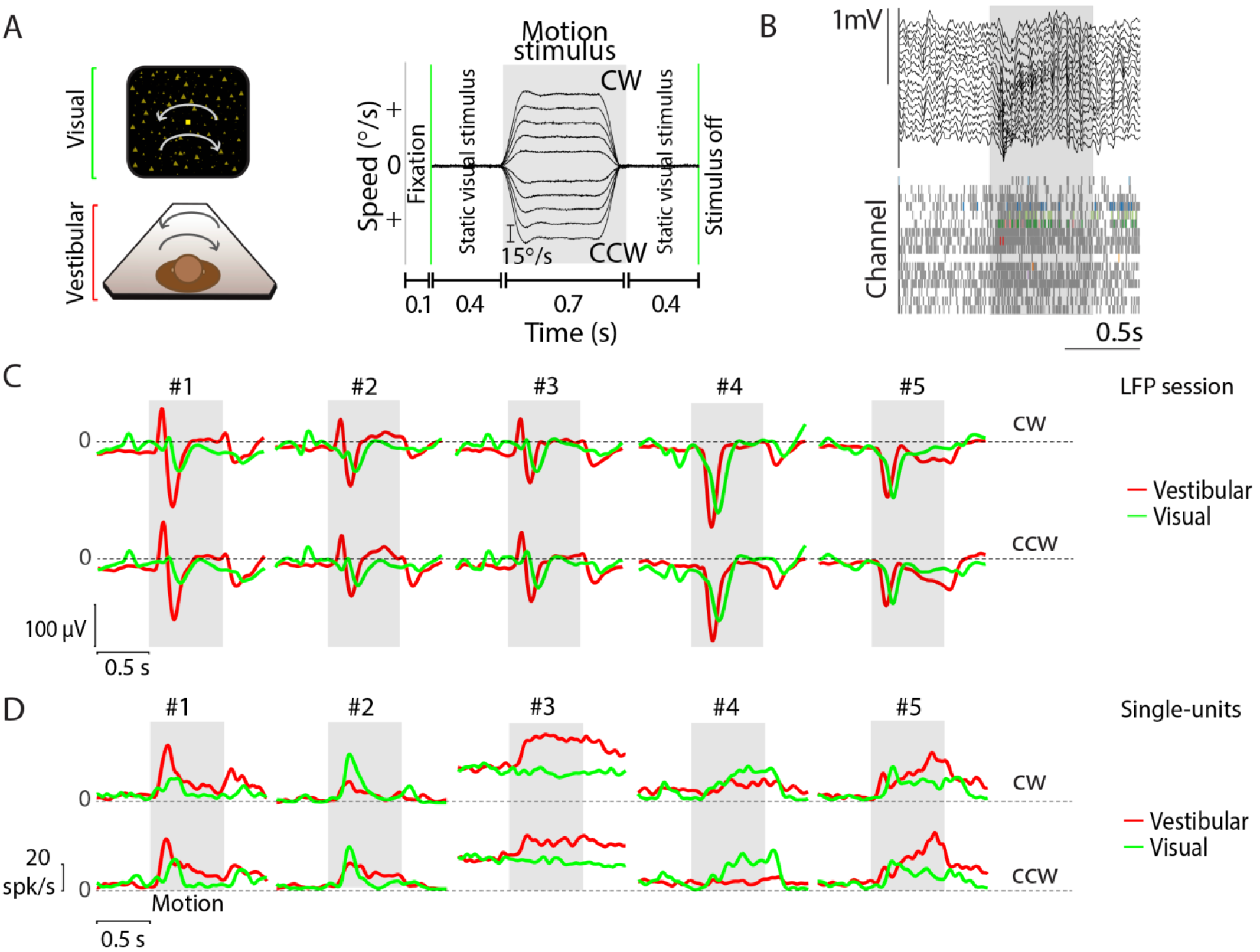
Example responses to angular rotation. **(A)** *Left-stimulus conditions’*. Monkeys were seated in a chair mounted on top of a rotary motor in front of a projector screen. Clockwise (CW) and counter-clockwise (CCW) rotational motion (yaw axis) was delivered visually via optic flow cues or by rotating the chair at various speeds. *Right-trial structure*: Time-course of rotational velocity reconstructed from accelerometer recordings obtained during ten different angular velocities (2 directions × 5 speeds) under the vestibular condition. Similar to linear translation, the desired temporal profile of angular velocity was trapezoidal for vestibular and visual rotation. Rotational velocity varied across trials from among ten possible values (+15°/s, +30°/s, +45°/s, +60°/s, and +75°/s). During trials with visual motion, the motion period was flanked by presentation of a static cloud of dots for 400 ms. The screen was blank throughout the trial under the vestibular condition. **(B)** Local field potentials (LFP) (top) and spikes (bottom, grey: multi-units, colored: single-units) recorded from all channels of the linear array during one example trial. **(C)** Stimulus-evoked LFPs (averaged across all channels) from five example recording sessions in response to vestibular (red) and visual (green) rotational motion for CW (top) and CCW (bottom) directions. **(D)** Time course of trial-averaged response of five example isolated neurons under the two conditions (visual and vestibular). Responses during trials with CW (top) and CCW (bottom) rotation were averaged separately across all speeds. Gray shaded areas show the motion period. CW: clockwise, CCW: counter-clockwise.

**Table 2.**
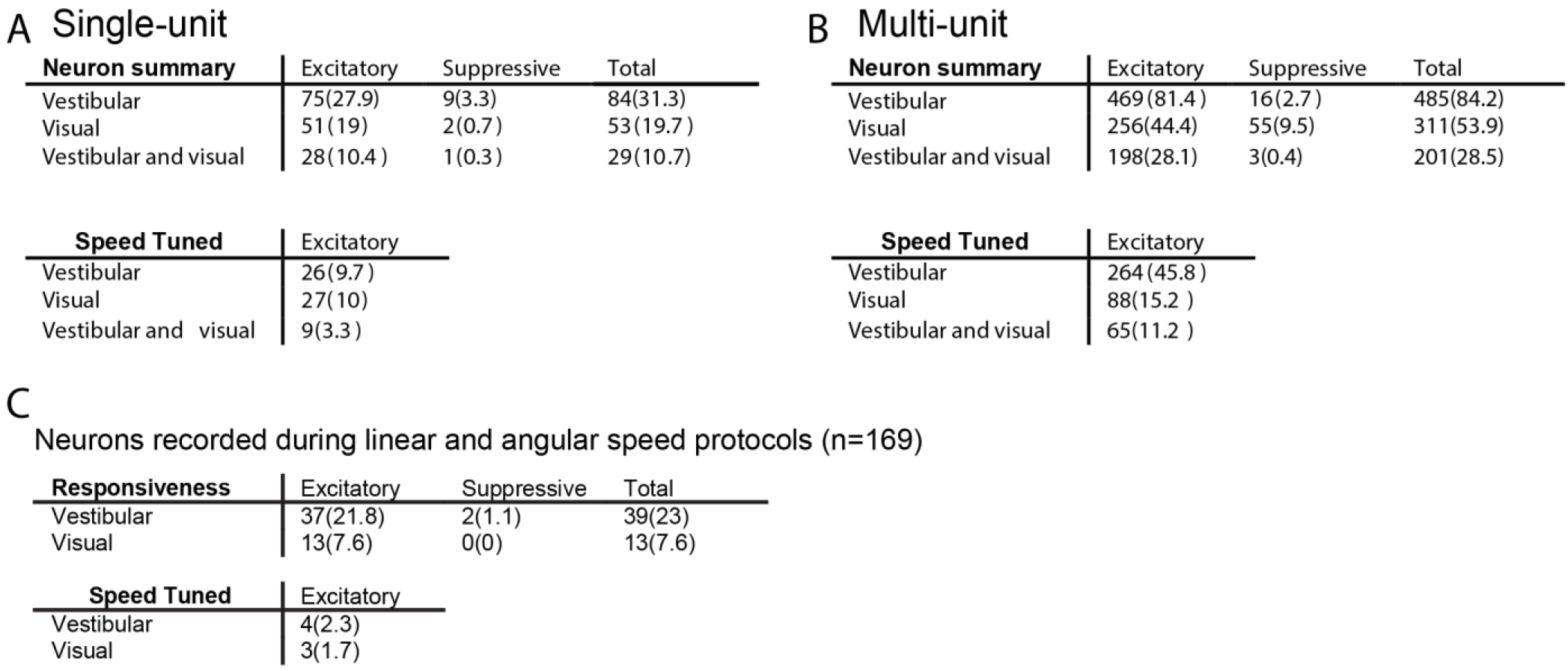
Statistics of neuronal and multi-unit response for angular speed. **(A)** Single-unit responses to angular rotation were classified in the same way as linear translation based on their response to the motion (excitatory, suppressive). **(B)** Corresponding statistics for multi-unit responses to angular rotation. **(C)** Statistics for 169 cells that were recorded under both linear and angular speeds. Values between brackets correspond to percentages of the total number of neurons recorded during the two protocols.

A Single-unit
| Neuron summary | Excitatory | Suppressive | Total |
| --- | --- | --- | --- |
| Vestibular | 75(27.9) | 9(3.3) | 84(31.3) |
| Visual | 51(19) | 2(0.7) | 53(19.7) |
| Vestibular and visual | 28(10.4) | 1(0.3) | 29(10.7) |

| Speed Tuned | Excitatory |
| --- | --- |
| Vestibular | 26(9.7) |
| Visual | 27(10) |
| Vestibular and visual | 9(3.3) |

B Multi-unit
| Neuron summary | Excitatory | Suppressive | Total |
| --- | --- | --- | --- |
| Vestibular | 469(81.4) | 16(2.7) | 485(84.2) |
| Visual | 256(44.4) | 55(9.5) | 311(53.9) |
| Vestibular and visual | 198(28.1) | 3(0.4) | 201(28.5) |

| Speed Tuned | Excitatory |
| --- | --- |
| Vestibular | 264(45.8) |
| Visual | 88(15.2) |
| Vestibular and visual | 65(11.2) |

C Neurons recorded during linear and angular speed protocols (n=169)
| Responsiveness | Excitatory | Suppressive | Total |
| --- | --- | --- | --- |
| Vestibular | 37(21.8) | 2(1.1) | 39(23) |
| Visual | 13(7.6) | 0(0) | 13(7.6) |

**Table 2. Statistics of neuronal and multi-unit response for angular speed.**
| Speed Tuned | Excitatory |
| --- | --- |
| Vestibular | 4(2.3) |
| Visual | 3(1.7) |

As was the case for linear translation, the effects described above were also present, and often much stronger, in multi-unit responses (**Supplementary Fig. S14, Table 2B**). Furthermore, across the population of all single-units, there was a significant difference in median SSI values between the vestibular (*SSI*=0.43 ±0.08) and visual (*SSI*=0.42±0.09) rotation conditions (paired *t*-test, *p*=0.036; **Fig. 5D, Supplementary Fig. S15**).

We wanted to know whether the same set of neurons were involved in encoding both linear and angular speeds. Of 169 single-units tested with a range of both translational and rotational speeds, ~23% of the neurons were responsive to vestibular motion and ~8% were responsive to visual motion under both protocols. Only ~2% of the units were significantly tuned to both linear and angular speeds (**Table 2C**), suggesting that linear and angular speeds are likely encoded by non-overlapping neural populations.

**Figure 5.**
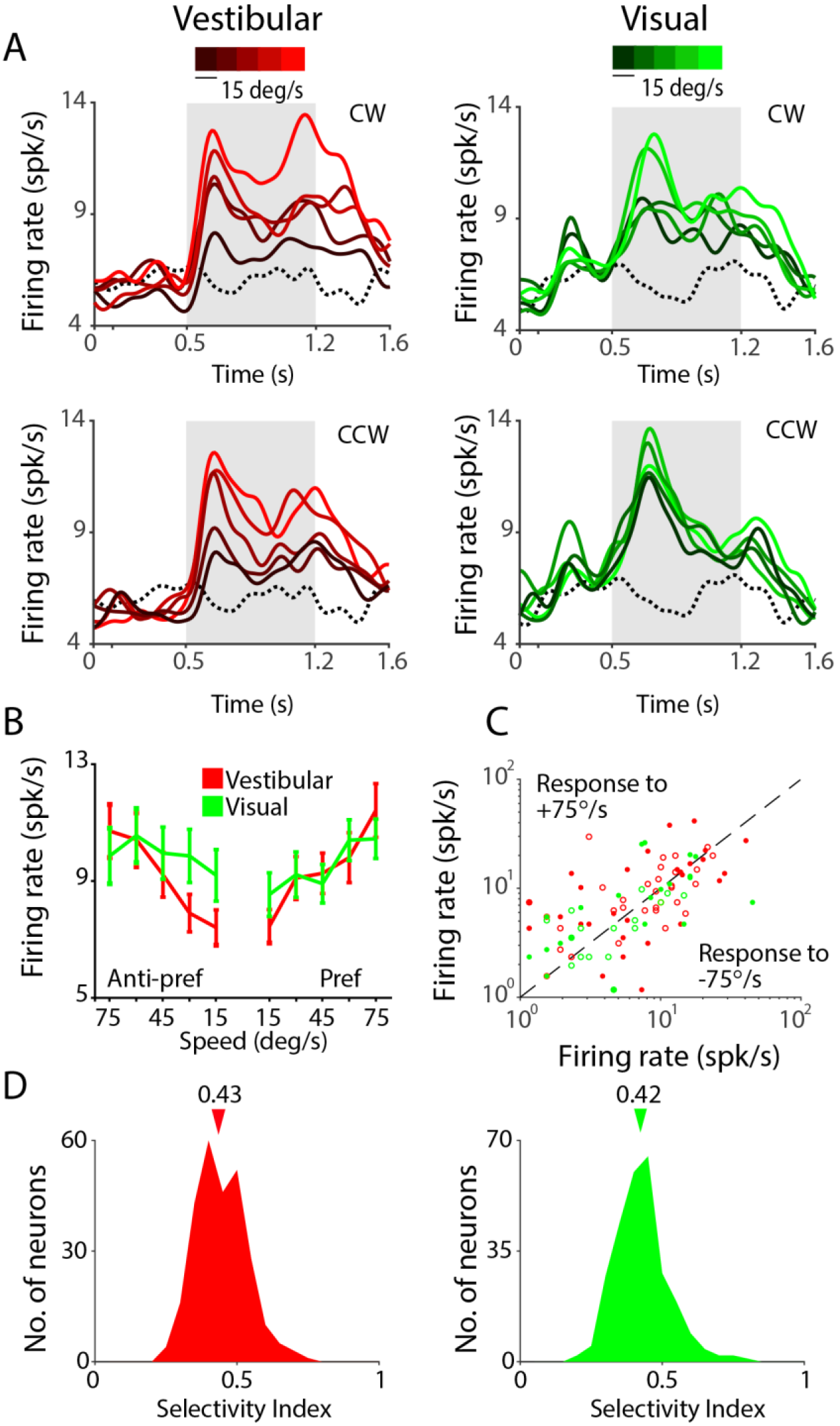
Selectivity to angular speed. **(A)** Time-course of response averaged across all neurons that were tuned to angular speed under vestibular (red, *n*=26) and visual (green, *n*=27) rotation conditions. Different shades correspond to different speeds (brighter hues correspond to greater speeds). Gray shaded areas show time-periods of motion. **(B)** Average speed tuning curves computed during the middle 500 ms period for the ‘preferred’ and ‘anti-preferred’ rotation directions (see text) **(C)** A comparison of the response of individual neurons to clockwise (+75°/s) and counterclockwise (75°/s) rotation. *open*- untuned, closed-tuned to angular speed under vestibular (red, *n*=55) or visual (green, *n*=25) conditions. **(D)** Distribution of the speed selectivity index (SSI) for all neurons in the two conditions. Arrows on top indicate medians of the respective distributions. CW: clockwise, CCW: counterclockwise.

### Spatial organization of responses

We leveraged the power of our laminar recordings, which were approximately normal to the cortical surface, to study the spatial organization of responses across the depth of a cortical column. We first tested whether similarities in the temporal fluctuations of LFP, single-unit, and multi-unit responses depended on the spatial separation of recording sites along a column. To test this, we concatenated temporal responses from all trials and computed pairwise correlations between the concatenated responses (this includes both signal and noise correlations). There was a clear distance-dependent decrease in shared variability of responses to both translational and rotational motion (**Fig. 6A**), which is well described by a power-law decay (**Supplementary Fig. S16**). Moreover, the correlation between LFP signals also exhibited a strong spatial periodicity with a wavelength of about 800 μm, suggesting a possible fine-scale structural organization of neural circuit processing in area 7a. For single-units and multi-units, a similar distance-dependent decrease was also found separately for correlations in stimulus-induced (signal correlation) and stimulus-independent (noise correlation) components of spike-count variability under both visual and vestibular motion conditions (**Supplementary Fig. S17**). These results suggest that neighboring neurons within a cortical column likely share similar response properties. Neurons in other regions of parietal cortex have previously been shown to be spatially clustered according to their heading tuning (Chen et al., 2008; Chen et al. 2010). We asked whether there was similar clustering of responses in area 7a based on speed selectivity. We found that the SSI values of single-units in response to both translational and rotational speed were more strongly correlated with those of multi-units recorded from the same electrode channel than those from other channels (**Fig. 6B, Supplementary Fig. S18**), suggesting that speed selective neurons are spatially clustered in 7a (though not necessarily in a columnar fashion).

**Figure 6.**
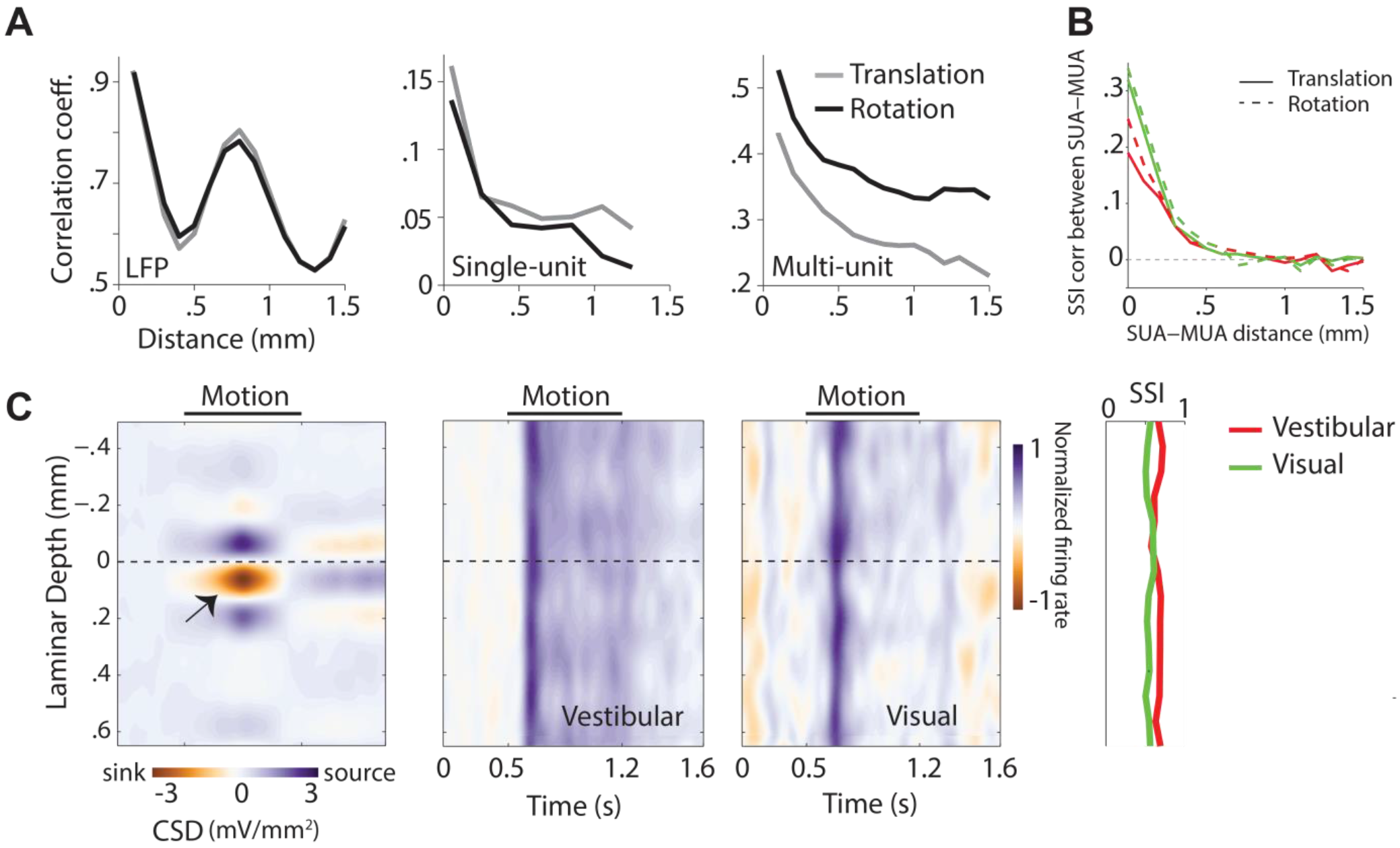
Spatial profile of responses. **(A)** Magnitude of correlated variability in the temporal fluctuations between pairs of simultaneously recorded local field potentials (*left*), single-unit spikes (*middle*), and multi-unit spikes (*right*), as a function of distance between the channels from which they were recorded. **(B)** Correlation in speed selectivity indices (SSI) between single- and multi-unit activity decreases as a function of the separation between the units. Red solid line: vestibular translation, green solid line: visual translation, red dashed line: vestibular rotation, green dashed line: visual rotation. **(C)** *Left:* Laminar profile of the current source density (CSD), averaged across recording sessions (*n*=14) that exhibited a clearly identifiable source and sink (indicated by black arrow). CSDs from individual recordings were shifted to align the bottom of putative layer IV (0mm, **Methods**) before averaging. *Right:* Average time course of the normalized multi-unit response to vestibular (left) and visual motion (right) as a function of cortical depth for all recording sessions (each session includes both protocols), aligned using the CSD as reference. The spatial profile of multi-unit SSIs for vestibular (red) and visual (green) motion is shown to the right. SUA: single-unit activity, MUA: multi-unit activity.

To test whether such clusters are confined to specific layers within a column, we used the laminar profile of the LFP to estimate the layers spanning the electrode array by measuring the current source density (CSD, **Methods**). While we cannot use CSD to tell the precise identity of the different layers, the location of the current sink can be used to identify putative layer IV and thereby the relative locations of the infra- and supra-granular layers (Schroeder, 1998; Maier et al., 2011; Self et al., 2013). We used this technique to align CSDs and multi-unit responses across different recordings (**Fig. 6C**). We found no systematic differences in SSI values across layers (one-way ANOVA, vestibular: *p*=0.41, visual: *p*=0.79). There was also no dependence of SSI on the coordinates of recording locations along the anterior-posterior (Translational SSI: Vestibular: Pearson’s correlation coefficient, *r*=0.03, *p*=0.72, Visual: *r*=0.05, *p*=0.57 ; Rotational SSI: Vestibular: *r*=−0.07, *p*=0.61, Visual: *r*=0.01, *p*=0.63), or the mediolateral axes (Translational SSI: Vestibular: *r*=0.07, *p*=0.49, Visual: *r*=0.11, *p*=0.37 ; Rotational SSI: Vestibular: *r*=−0.06, *p*=0.52, Visual: *r*=0.15, *p*=0.51). Together, these results suggest that speed tuning is not restricted to specific layers or sites in area 7a (**Supplementary Fig. S19**).

### Precision of the speed code

The speed-selective neurons reported here are likely to play a fundamental role in path integration, the process of computing one’s position by integrating self-motion velocity signals. We evaluated the precision of translational and rotational speed representations in our data to test how well a hypothetical decoder of the recorded neural activity can estimate travel distances and angles (**Fig. 7A**). We made the following assumptions in order to do this. First, most neurons that were sensitive to rotational speed did not discriminate between the two directions of rotation. Since self-motion under natural conditions is mostly in the forward direction, we did not test neuronal sensitivity to translational speed under backward heading. Therefore, we do not know if neurons in area 7a encode the direction of linear translation. We assume that accurate information about the direction of translation (forward/backward) and rotation (clockwise/counterclockwise) is available to the velocity decoder, presumably from other areas such as MSTd (Takahashi et al., 2007; Yang et al., 2011) that encode those quantities. Second, we assume that the encoding of speed is instantaneous so any temporal filtering in responses induced by cellular or synaptic mechanisms will not significantly alter the precision of the code under naturalistic conditions involving continuous movements. Finally, based on evidence from human behavioral studies (Lakshminarasimhan et al., 2017, preprint), we assume that brain mechanisms that integrate self-motion velocity are nearly perfect so that errors in estimating movement distances and angles are largely due to uncertainty in the decoded velocity estimates. Under these assumptions, we can examine the neural representation of speed to predict errors in estimating distances and angles during path integration.

**Figure 7.**
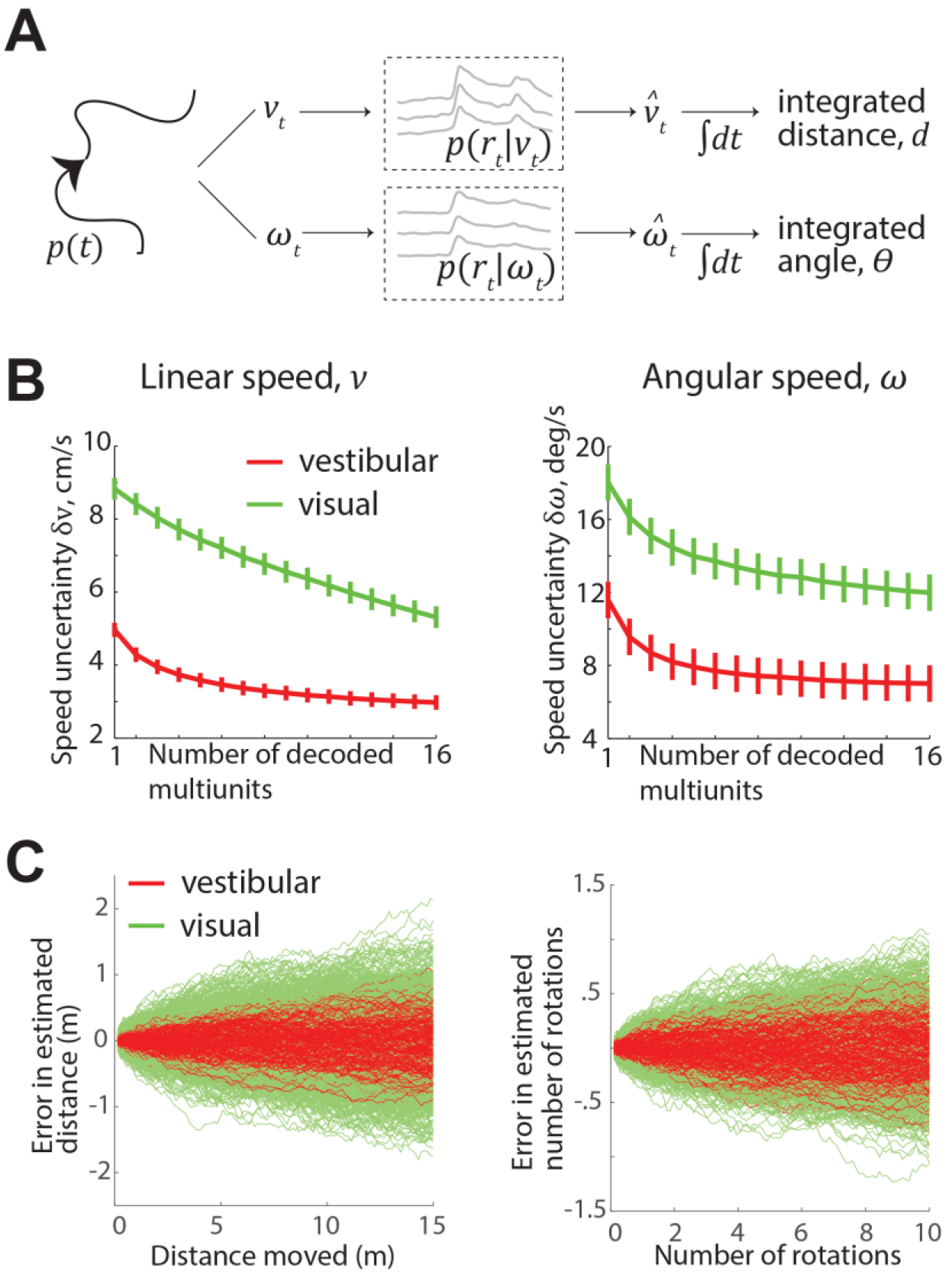
Precision of the speed code. **(A)** A diagram illustrating the role of speed-selective neurons in path integration. Any generic trajectory *p(t*) can be decomposed into time-varying translational (*v_t_*) and rotational (*ω_t_*) velocities that can be encoded in the activity of neurons, as characterized by the response distributions *p*(*r*|*v*) and *p*(*r*|*ω*), respectively. We can decode the joint activity of these neurons to generate estimates of translational and rotational velocity 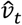 and 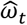), which can then be integrated to compute distance moved and angle turned respectively. (*B*) Uncertainty in estimates of translation (*δv, left*) and rotation (*δω*, *right*) speeds generated by decoding responses along the principal component of the multi-unit response distributions *p*(*r*| *v*) and *p*(*r*|*ω*). These uncertainties are plotted as a function of the number of simultaneously recorded multi-units used to decode speed. Error bars denote ±1 standard error of the mean (across different recording sessions (*n*=44 and *n*=36 for linear and angular respectively) value of uncertainty. **(C)** *Left:* Simulated errors in estimated distance by integrating decoded linear speed, plotted as a function of the actual distance moved under vestibular (red) and visual (green) conditions. *Right*: Errors in estimating angle (1 rotation = 360°) as a function of the true angle. Results from 1000 simulations overlaid for each case.

Uncertainty in the estimated velocity depends on the choice of the decoder. For linear population codes, a decoder with minimal uncertainty will be one that takes the fine structure of correlated variability into account. However, this fine structure is difficult to assess experimentally with limited data so it is practically impossible to determine the optimal decoder-one that will yield estimates with least uncertainty. Instead, we considered a decoder that extracts velocity information available within the principal component of the population response. The uncertainty implied by this decoder is simply the inverse square root of the signal-to-noise ratio along the principal component of the population response. Here, signal and noise correspond to the slope of the tuning curves and variance of the neural activity projected into the principal component subspace respectively (**Methods**). We computed the uncertainty by considering ensembles of 1-16 simultaneously recorded (single session) multi-units and by averaging across many recordings separately for the vestibular and visual conditions of translational and rotational motion (**Fig. 7B**). As expected from the response properties of single neurons detailed in the previous sections, uncertainties of speed estimates were smaller for the decoder based on vestibular responses than visual responses (vestibular: *δv* = 3.1±0.4cm/s and *δω* = 7.8±2.1°/s; visual: *δv* = 4.9±0.7cm/s and *δω* = 13.2±2.3°/s for the largest ensemble size, *n*=16 at *v*=13.75cm/s and *ω*=37.5°/s).

Uncertainty in velocity estimates would build up to produce errors in path integration. To evaluate the performance of the decoder in path integration, we integrated the output of this decoder to estimate the distance moved and angle turned under conditions of pure translation (at a speed of 13.75 cm/s) and pure rotation (at 37.5 °/s), respectively. These speeds were chosen as they corresponded to the mean speeds tested in our experiments. Integrated distance and angle estimates were generated according to a random walk by accumulating temporally independent samples drawn from a normal distribution with unbiased means (equal to the actual speeds) and standard deviation equal to the uncertainty resulting from decoding linear and angular speeds using the largest ensemble size (*n*=16). This procedure was repeated many times to obtain average estimation errors in distance (**Fig. 7C**-*left*) and angle (**Fig. 7C**-*right*). Estimates based on visual cues were much worse as the uncertainties in both translational and rotational speed estimates generated by decoding visual responses were greater than those generated by decoding vestibular responses. Nevertheless, both visual and vestibular estimates of distance and angle were found to be well within 10% of the actual values suggesting that an ideal integrator of the activity of neurons in area 7a can, in principle, maintain qualitatively good estimates of position based solely on self-motion cues (mean estimated distance at 15m-visual: 15 ± 0.5 m, vestibular: 15 ± 0.3 m; mean estimated number of rotations after 10 rotations-visual: 10 ± 0.35, vestibular: 10 ± 0.2).

We emphasize that decoding velocities along the principal component represents a very conservative choice because this component represents the noisiest dimension of neural responses and will inflate the expected uncertainty. Moreover, although there was a hint of saturation in our estimates of uncertainty in decoding speed, decoding larger ensembles will lead to smaller errors. Notwithstanding potential errors arising from any additional noise in the integration mechanism and uncertainty in direction estimates, we believe that the above error estimates likely represent a theoretical upper bound on path integration errors.

## DISCUSSION

We have found evidence for a multisensory representation of self-motion velocity in area 7a of the macaque PPC. Our findings extend the literature on self-motion processing in this brain area in three ways. First, past studies on speed sensitivity of neurons in area 7a used radial optic flow stimuli typically experienced during fore-aft translation (Siegel and Read, 1997; Phinney and Siegel, 2000), but did not examine responses to optic flow that simulates angular rotation. Here we used expanding as well as rotational flow patterns to investigate the representation of both translational and rotational components of self-motion velocity. Second, to our knowledge, sensitivity of area 7a neurons to vestibular inputs has never been explicitly tested. A previous study by Kawano et al. 1980 reported vestibular responses from area 7. However, these recordings were in fact performed in the anterior bank of the superior temporal sulcus (area MST, (Kawano et al., 1980; Komatsu and Wurtz, 1988), K. Kawano, personal communication, 2016). Here, we used a motion platform with translational and rotational degrees of freedom to explore representations of selfmotion velocity based on vestibular cues. Finally, for the first time we used combined visual-vestibular stimuli to study multisensory convergence of self-motion cues during forward translation in area 7a.

Our experiments using visual motion stimuli confirm earlier reports of optic flow sensitive neurons in area 7a (Siegel and Read, 1997; Read and Siegel, 1997; Merchant et al., 2001; Raffi and Siegel, 2007). The fraction of neurons with a statistically significant response to visual translation in our data was roughly similar to those studies for the conditions in which flow fields were similar (~30% for Siegel and Read, 1997, and Merchant et al. 2001). The overall greater percentage of flow-responsive neurons in these previous studies is likely because of the greater variety of optic flow patterns that were employed. For example, Merchant et al. (2001) found that ~60% of neurons in area 7a responded significantly to at least one of their flow fields. In our recordings, we limited the optic flow conditions to those corresponding to natural navigation, that is, expanding flow (straightforward linear translation) and rotational flow (yaw rotation) only, which limited the fraction of neurons that could be driven effectively by our stimuli. In addition, recordings in previous studies were carried out using single electrodes, which may allow for greater sampling biases in selecting neurons. In contrast, we used linear electrode arrays in order to increase yield and we isolated all single-units offline before assessing their responsiveness, thus reducing sampling bias. The fraction of visually responsive multi-units in our dataset was substantially greater than for single-units, and was more comparable to earlier work using a greater variety of optic flow (Siegel and Read, 1997; Read and Siegel, 1997; Phinney and Siegel, 2000; Merchant et al. 2001). In addition, we observed significant motion-induced (negative) evoked potentials in the LFP for the vast majority of recording sites, implying widespread activation of area 7a by optic flow. The sign, shape and amplitude of the LFP depends on the recording position, morphology and position of the synapses (Einevoll, 2013). On average, synaptic excitation will cause currents to leave the extracellular region (and enter the neurons) leading to a decrease in potential in the extracellular space. Hence, the negative deflections in the LFP caused by the stimulus (Fig. 1C, Supplementary Fig. 2A).

A previous study on speed selectivity for optic flow reported heterogeneous tuning to speed with individual 7a neurons responding exclusively to low, high, intermediate, or multiple different speeds (Phinney and Siegel, 2000). We did not observe such diversity in speed tuning—responses of most speed selective neurons in our recordings increased with speed. Notably, earlier work on area MSTd using radial optic flow under conditions similar to ours also found mostly monotonically increasing speed tuning (Duffy & Wurtz, 1997). The differences between our findings and those of Phinney and Siegel (2000) are likely due to some fundamental differences between the two studies. First, Phinney and Siegel (2000) used a wider range of speeds that we did, which may have led to greater diversity in speed tuning profiles. Second, Phinney and Siegel (2000) used a wider variety of optic flow patterns including contraction, spirals, and linear translation along various directions on the horizontal plane. It is possible that speed tuning is non-monotonic for some optic flow patterns. Third, their recordings were performed in behaving monkeys that were trained to detect changes in the global structure of optic flow, not speed. Sensory representations in the PPC are modulated by attentional factors and continuously refined during learning, and therefore sensitive to task demands (Bucci, 2009; Robinson and Bucci, 2012). Consequently, the nature of speed tuning might also have been influenced by the task that Phinney and Siegel (2000) used.

Our experiments using the motion platform revealed that neurons in area 7a are sensitive to both translational and rotational vestibular cues. There is previous indirect evidence of a vestibular influence on neuronal responses in 7a (Brotchie et al., 1995; Snyder et al., 1998). Specifically, head orientation was found to modulate visual responses only when the head rotates above the vestibular threshold for a particular orientation. Here, we demonstrate a direct effect of physical movement (both translation and rotation) on the responses of individual 7a neurons, as well as MUA. Critically, we find that many 7a neurons signal the speed of vestibular translation and/or rotation, indicating that area 7a can use vestibular inputs to convey explicit information about self-motion speed, not just head orientation (Snyder et al., 1998). The origin of vestibular inputs to area 7a may be either from connections with thalamic nuclei that convey vestibular input to the cortex (Ventre and Faugier-Grimaud, 1989), or from multimodal areas that project to 7a, such as MSTd and the ventral intraparietal (VIP) area (Pandya and Seltzer, 1982; Rozzi et al., 2006; Seltzer and Pandya, 1986; Van Essen et al., 1990). Additionally, it is worth noting that area 7a may project directly to the vestibular nuclei (Faugier-Grimaud and Ventre, 1989).

Responses to the combined visual-vestibular stimulus tended to be dominated by vestibular, rather than visual, inputs (**Supplementary Figs. S4, S7B**). In some cases, neurons selectively responsive to visual motion were even suppressed when visual motion was paired with platform movement (Fig. 1D #2). This dominance of vestibular influences on responses was unexpected because area 7a is thought to be largely visual and is generally considered the output of the dorsal visual hierarchy (Andersen, 1989; Britten, 2008). Moreover, vestibular dominance has not been reported in other multimodal parietal areas, such as MSTd and VIP (Schlack et al., 2002; Bremmer et al., 2002(b); Gu et al., 2008; Chen et al., 2013). The fact that vestibular inputs more strongly dictate neuronal responses in area 7a suggests that local mechanisms may inhibit visual motion inputs when vestibular cues are available. Interestingly, a circuit mechanism involving parvalbumin-expressing interneurons in the mouse PPC has recently been found to help resolve multisensory conflicts in audiovisual tasks (Song et al., 2017). It is possible that area 7a could perform an analogous role in mediating conflicts related to self-motion processing in macaques, although this remains a speculation at this point. Such a role would be consistent with the idea that this area conveys self-motion information to navigation circuits, for it is critical to resolve such conflicts before using self-motion to navigate.

Across both conditions, neuronal responses often varied with speed. This result parallels the outcome of experiments in freely moving rodents that demonstrate speed representation by neurons in the PPC (Whitlock et al., 2012). However, unlike in rodent PPC, neurons in area 7a did not show diverse preferences for speed. Instead, their responses were more similar to ‘speed cells’ found in the medial entorhinal cortex of rats that exhibit a speed-dependent increase in firing rate (Sargolini et al., 2006; Kropff et al., 2015; Hinman et al., 2016). Our recordings also revealed an important qualitative feature of angular velocity representation in 7a-firing rates increased with rotation speed regardless of the direction of yaw rotation. Thus, while the magnitude of angular speed could be decoded from this area, information about rotation direction (clockwise or counter-clockwise) likely has to be obtained from other brain areas. Although speed-dependent increases in firing rate were found for both translation and rotation, neuronal populations representing translational and rotational speeds were almost completely non-overlapping (Table 2C). A potential computational benefit of such a spatially de-multiplexed representation is that the translational and rotational components may be decoded independently by reading out the corresponding populations, and integrated separately to estimate distance and heading.

A decrease in temporal response correlations with distance (Supplementary Figs. 16, 17) supports the notion that neurons are spatially clustered with respect to their sensory representation of motion. Moreover, the heterogeneity in responses across sites suggests that there is no clear topographic organization for a specific modality or for the strength of speed tuning (Supplementary Fig. 19). Most of the recording penetrations were approximately normal to the surface of the cortex, such that the clustering is within the span of a column although we cannot rule out small variations in the penetration angle of the recordings. Functional organization of speed tuned neurons has been described in macaque area MT (Liu and Newsome, 2002) where penetrations normal to the surface of the cortex lack a columnar organization and rather show neurons clustered according to preferred speed. Similarly, they also observe that single and multi-unit activities recorded from the same sites were correlated. Spatial clustering could be a result of anatomical connections. Overlapping connections have been described in the inferior parietal lobule where prefrontal and parietal fibers overlap in area MST (Seltzer et al., 1996). Moreover, a recent study in mice by Yamawaki et al. (2016) describe two projections from different areas to PPC in which the afferents are interspersed. In addition to coarse clustering of neuronal responses, a finer scale in the organization of the responses may also be present. In fact, we observed that the LFPs exhibit spatial periodicity. We used CSD to test if there was a layer-specific encoding of speed. Unlike other areas (Self et al., 2013; Nandy et al., 2017), we found no layer-specific differences in the strength of tuning or multi-unit response latencies. This is consistent with recent findings from the rat parietal cortex showing self-motion tuned neurons are found across all depths (Wilber et al., 2017).

Precise knowledge of self-motion velocity is indispensable for path integration. We quantified the precision of speed representation in area 7a by testing the performance of a hypothetical decoder estimating distances and angles. Unlike approaches that use advanced decoders based on suppor*t*-vector machines (Liu et al., 2012), logistic regression (Berens et al., 2012), or likelihood-based **Methods** (Graf et al., 2011), we considered a simple readout along the principal component of the population response due to the limited numbers of trials in our data sets. Yet, our conservative readout scheme yielded distance and angle estimates that were well below the behavioral precision of humans and other animals performing path integration (Glasauer et al., 1994; Etienne et al., 1996; Israël et al., 1997; Maaswinkel and Whishaw, 1999). This supports the idea that the speed representation in area 7a is, in principle, sufficient to support integration over a modest range of translation and rotation. Further experiments involving inactivation and/or perturbation of neural activity in area 7a will be required to establish a causal link to path integration behavior. Imaging studies in humans have identified candidate areas of the PPC involved in navigation and route planning using virtual reality tasks (Maguire, 1998; Rosenbaum et al., 2004; Shelton and Gabrieli, 2002; Wolbers et al., 2004; Spiers and Maguire, 2006, 2007; Ciaramelli et al., 2010). Similar tasks in macaque monkeys have revealed route-selective neurons in the medial parietal region (Sato et al., 2006, 2010). Our findings complement those studies by demonstrating a potential role for the PPC in self-motion-based, rather than route-based, navigation.

Although we focused on multisensory coding of self-motion information in this work, the PPC could span multiple levels of a hierarchy of representations (Chafee and Crowe, 2012). In this general framework, single neurons in PPC may carry mixtures of signals at many different time scales, ranging from sensory variables such as movement speeds reported here to dynamical representations that are modulated by working memory (Constantinidis & Steinmetz, 1996; Qi et al., 2010; Rawley & Constantinidis, 2010), attention (Steinmetz et al., 1994; Robinson et al., 1995; Constantinidis & Steinmetz, 2001a,b; Quraishi et al., 2007; Oleksiak et al., 2011), or learning (Freedman & Assad, 2006). The flow of information between brain areas is likely modulated by task demands, which could in turn influence precisely where neuronal responses fall along this continuum. We know relatively little about the neural mechanisms by which interactive environments can dynamically lead to the emergence of cognitive neural signals in the cortex. Future studies should use complex, naturalistic tasks and normative models of behavior to understand the flow of information between populations of neurons that encode different task-relevant variables, and how the PPC transforms sensory inputs into a more behaviorally relevant format.

## METHODS

### Animal preparation

We recorded extracellularly using linear array electrodes from area 7a of three adult male rhesus macaques (*Macaca Mulatta*; 8.5-10 kg). We collected data from 9 recording sessions from the left hemisphere and 10 from the right hemisphere from Monkey O, 5 sessions from the right hemisphere of monkey J, and 20 sessions from the right hemisphere of monkey Q. All animals were chronically implanted with a lightweight polyacetal ring for head restraint, a removable grid to guide electrode penetrations, and scleral coils for monitoring eye movements (CNC Engineering, Seattle WA, USA). Monkeys were trained using standard operant conditioning procedures to maintain fixation on a visual target within a two degree square window for a period up to three seconds. All surgeries and procedures were approved by the Institutional Animal Care and Use Committee at Baylor College of Medicine, and were in accordance with National Institutes of Health guidelines.

### Experimental setup

Monkeys were head-fixed and secured in a primate chair that was mounted on top of a rotary motor (Kollmorgen, Radford, VA, USA). The chair and motor were all mounted on a 6 degree-of-freedom motion platform (Moog 6DOF2000E, East Aurora, NY, USA) that was used to deliver two different vestibular stimulus protocols (see below). A 3-chip DLP projector (Christie Digital Mirage 2000, Cypress, CA, USA) was mounted on top of the motion platform and rear-projected images onto a 60 × 60 cm tangent screen that was attached to the front of the field coil frame, ~30 cm in front of the monkey. The projector was capable of rendering stereoscopic images generated by an OpenGL accelerator board (Nvidia Quadro FX 3000G). The display was updated at 60 Hz, synchronous with commands to update the position of the motion platform. Custom scripts were written for behavioral control and data acquisition with TEMPO software (Reflective Computing, St. Louis, MO, USA).

### Stimulus

Monkeys had to maintain fixation on a central fixation target (a square that subtended 0.5 deg on each side) for 100 ms to initiate each trial. A self-motion stimulus that lasted 700 ms was presented during each trial. The onset of motion occurred 400 ms following trial initiation, and its dynamics followed a trapezoidal profile with the speed held fixed for 500 ms in the middle (**Fig. 1A**). Depending on the experimental protocol (see below), up to three different kinds of trials were interleaved in which the self-motion stimulus was presented by moving the platform/chair in which the animal was seated (vestibular condition), by presenting visual motion of a 3-dimensional cloud of dots having a density of 0.01 dots/cm^3^ (visual condition), or through a congruent combination of both cues (combined condition). In trials containing visual motion, presentation of the motion stimulus was flanked on either side by a 400 ms display of a static cloud of dots to prevent on/off visual transient responses from contaminating neural responses to motion. In the vestibular condition, no visual stimulus was present during these periods, except for the fixation square. A liquid reward was delivered at the end of the trial if the monkey maintained fixation throughout the trial. Trials were aborted following fixation breaks.

In each recording session, we employed two different experimental protocols-*translation* and *rotation*- that shared a trial structure identical to that described above but differed in the type of motion presented: whereas the former involved linear translation along the forward (straight-ahead) direction, the latter involved a pure yaw rotation. Vestibular presentation of translational motion was achieved by translating the motion platform, while presentation of rotational motion involved rotating the motor beneath the monkey’s chair. Note that the projection screen was attached with to the motion platform and translated with the animal, whereas yaw rotations applied by the motor changed the animal’s body orientation relative to the projection screen. Whereas the translation protocol included all three stimulus modality conditions (visual/vestibular/combined), the rotation protocol included only the two unimodal conditions as combined presentation of the rotation stimulus was technically challenging. For both the translation and rotation protocols, the visual fixation target remained head-fixed during movement, such that no eye-in-head movements were necessary for the animal to maintain fixation.

The speed of motion was varied across trials in each protocol. We employed four different speeds of linear translation between 5.5 and 22cm/s in steps of 5.5cm/s, and five different rotation speeds (15-75°/s in steps of 15°/s) for both clockwise and counter-clockwise directions of rotation. We collected data for at least five repetitions of each distinct stimulus, and both protocols included additional control trials (baseline condition) in which the monkey was required to fixate on a blank screen.

### Recording and Data acquisition

Area 7a was identified using structural MRI, MRI atlases, and confirmed physiologically with the presence of large receptive fields responsive to optic flow (Siegel and Read, 1997; Merchant et al., 2001). Extracellular activity was recorded using a 16-channel linear electrode array with electrodes spaced 100 apart (U-Probe, Plexon Inc, Dallas, TX, USA). The electrode array was advanced into the cortex through a guide-tube using a hydraulic microdrive. The spike-detection threshold was manually adjusted separately for each channel to facilitate monitoring of action potential waveforms in real-time, and recordings began once the waveforms were stable. The broadband signals were amplified and digitized at 20 KHz using a multichannel data acquisition system (Plexon Inc, Dallas, TX, USA) and were stored along with the action potential waveforms for offline analysis. Additionally, for each channel, we also stored low-pass filtered (-3dB at 250Hz) local-field potential (LFP) signals.

### Recording reconstructions

Two MRI data sets of T1-weighted scans from monkey Q were used to reconstruct recording sites using a frameless stereotaxic technique pre and post ring implantation. The latter included a plastic grid filled with betadine ointment to allow for precise electrode targeting and reconstruction of recordings sites. We used an operator-assisted segmentation tool for volumetric imaging (ITK-SNAP v3.6, www.itksnap.org) to segment in 3D and render the brain structures from the MRI scans (Yushkevich et al., 2006). We aligned the MRI datasets to bony landmarks and brain structures to identify anterior-posterior zero (ear bar zero). All horizontal slices were aligned to the horizontal plane passing through the interaural line and the infraorbital ridge; all coronal slices were aligned parallel to the vertical plane passing through the interaural line (ear-bar zero). Brain structures used for alignment were the anterior and posterior commissure, as well as the genu and splenium of the corpus callosum (Dubowitz and Scadeng, 2011; Frey et al., 2011). All of the anatomical landmarks derived from horizontal, coronal, and sagittal sections were then used to index to positions in standard macaque brain atlases (Paxinos et al., 2000; Saleem and Logothetis, 2012) and recording sites were then reconstructed using the position on the grid in stereotaxic coordinates.

### Data analysis

Spikes were sorted offline using commercially available software (Offline Sorter v3.3.5, Plexon Inc, Dallas, TX, USA) to isolate single-units. Spikes that could not be distinctly clustered were grouped and labeled as multi-unit activity (MUA), representing a measure of spiking of neurons in the vicinity of the electrode. All subsequent analyses were performed using Matlab (The Mathworks, Natick, MA, USA). Stimulus-evoked local field potentials (LFP) in **Figures 1** and **4** were obtained by trial-averaging LFP traces recorded from each electrode channel, and then averaging across electrodes. Similarly, for single-units and MUA, we constructed peri-stimulus time histograms (PSTHs) for the complete duration of each trial (1.6 s including the fixation period) in each stimulus condition by convolving the spike trains with a 25 ms wide Gaussian function and then averaging across stimulus repetitions.

A neuron was classified as responsive to motion if its average firing rate during a 400 ms time-window following motion stimulus onset was significantly different from its activity during a window of the same length preceding motion onset (*p* ≤ 0.05; two-sided *t*-test). Responsive neurons were further classified into “excitatory” and “suppressive” depending on whether motion increased or decreased neuronal activity relative to baseline. We tested the significance of tuning for speed by performing a one-way ANOVA (*p* ≤ 0.05) to test if the mean responses to different speeds were significantly different from each other. We estimated tuning functions for speed by measuring the average firing rate during the 500 ms flat phase of the trapezoidal motion profile when speed was held constant. To quantify the strength of tuning to speed, we computed a speed selectivity index (*SSI*) which ranged from 0 to 1 (Takahashi et al. 2007): 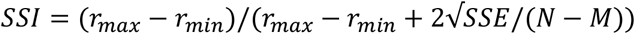 where *r*_*max*_ and *r*_*min*_ are the mean firing rates of the neuron in response to speeds that elicited maximal and minimal responses, respectively; SSE is the sum-squared error around the mean responses; *N* is the total number of observations (trials); and *M* is the total number of speeds (4 and 5 for the translation and rotation protocols respectively). For rotational motion, we averaged responses to the two rotation directions separately for each rotation speed before computing *r*_*max*_ and *r*_*min*_.

For LFPs, peak-amplitude was defined as the amplitude of the largest peak or trough in the signal after motion onset. For both spikes and LFP, response latency was defined as the time after the onset of motion at which the activity deviated by 2 standard deviations from the average response during the 400 ms window prior to motion onset, for at least four consecutive time bins (100 ms total). This specific comparison with the time window preceding motion onset was chosen to control for responses related to the appearance of static dots at the onset of the visual stimulus. We also analyzed the temporal dynamics of single- and multi-unit activity to determine whether the temporal response was unimodal or bimodal. To do this, we determined the time-points at which the firing rate within a 25 ms sliding window was at least 2 standard deviations above the response in the immediately preceding window. We identified such-time points within the 400 ms window following motion onset and offset (if available) to categorize single- and multi-units into unimodal and bimodal units. Units with at least one such time point following motion onset, but none following motion offset, were classified as unimodal, whereas neurons with such time points following both onset and offset of motion were labelled bimodal.

To compute the correlation between neural responses and motion speed, we computed the trial-averaged response of individual neurons to each possible speed. We then pooled the trial-averaged responses of all neurons to obtain a vector of responses to each motion speed, and estimated the correlation between the resulting population vectors and motion speed.

### Pairwise correlations

We computed temporal correlations between responses of simultaneously recorded pairs of single-units, multi-units, and LFPs. For single- and multi-units, we first smoothed spike trains from individual trials using a 25ms Gaussian window before concatenating temporal responses of all trials. For LFPs, we concatenated the raw voltage signals recorded during each trial. Finally, we computed pairwise correlations between the resulting smoothed concatenated time-series.

For single- and multi-units, we additionally computed signal and noise correlations separately for vestibular and visual conditions. Signal correlation was computed as the Pearson correlation between the speed tuning curves (obtained from the average firing rate during the middle 500ms of the trapezoidal motion) of pairs of units. For computing noise correlations, we z-scored average firing rates (during the middle 500ms of the trapezoidal motion) separately for trials corresponding to each motion speed, and then computed pairwise correlations between pooled z-scored responses. The z-scoring essentially removes shared fluctuations induced by changes in speed (signal), but retains correlations in trial-by-trial variability (noise). We estimated pairwise correlations as a function of distance by pooling all pairs with a given electrode separation (0 to 1500μm in steps of 100μm), regardless of depth.

Current Source Density (CSD), *J*, was computed as the second spatial derivative of the LFP *Ψ* according to *J*(*x, t*) = *∂*^2^*Ψ*(*x, t*)/*∂x*^2^, which we discretized as *J*(*i, t*) = (2*Ψ*(*i, t*) − *Ψ*(*i* + 1,*t*) − *Ψ*(*i* — 1, *t*))/(Δ*x*)^2^. Here *i* and *t* correspond to the channel number and time respectively, and Δ*x* = 0.1 *mm* was the inter-channel spacing between our electrode sites. CSD was computed using 3 s long LFP segments averaged across trials. Following standard conventions, sites with positive and negative densities were labelled as sources and sinks, respectively. To align CSD and multi-unit responses from different recordings, we computed the centroid of the sink at the time-point where the current density was most negative. CSD and multi-unit responses from different recordings were shifted across space to align the respective centroids before averaging. The top of the sink was labelled as the bottom of putative layer IV following standard convention used for labeling layers in macaque primary visual cortex (Mitzdorf and Singer, 1979; Schroeder, 1998; Self et al., 2013).

### Spike waveform classification

Neurons were classified into putative pyramidal and interneurons using standard **methods** based on the shape of the action potential waveform (Mitchell et al., 2007). We constructed the distribution of spike waveform half-widths (defined as the average peak-trough duration). We used a Hartigan’s dip test (*p*<0.05, Hartigan and Hartigan, 1985) to test for bimodality and neurons were categorized into narrow and broad spiking depending on the percentile score of their spike half-widths. Where necessary, we inverted the polarity of spike waveforms to ensure that peaks followed troughs. We excluded 11 neurons from this analysis as the peaks and troughs of the action potential could not be reliably determined.

### Linear Decoder

Covariance ∑ was computed for each ensemble of simultaneously recorded units by combining z-scored responses from all trials (similar to the method described above for computing noise correlations). In principle, the uncertainty *δ* in the neural representation of speed is given by the square-root of the Cramer-Rao bound, which is equal to the inverse square-root of the linear Fisher information, *I*, according to 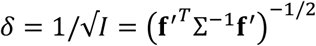 where **f**′ is the vector of derivatives of the neuronal tuning curves around the mean estimate (the actual speed). However, in practice, it is very difficult to reliably estimate the covariance ∑ with limited data, so the Cramer-Rao bound cannot be used. One can still reliably estimate the leading modes of covariance (leading principal components) as they have the largest
variance. Therefore, we considered a simple readout scheme in which the weights of the linear decoder were aligned along the leading principle component **u** of the population response covariance ∑. If *λ* is the variance of the neural response along the principal component **u**, then the linear Fisher information along this direction is equal to 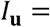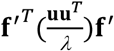, and the uncertainty is given by 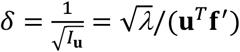. Uncertainties in linear and angular speeds were estimated separately from ensembles of simultaneously recorded multi-units (*n*=1 to 16) within each session. We used the average uncertainty across sessions for the largest ensemble (*n*=16) as the uncertainty of the decoder. To determine theoretical errors in estimating distance and angle from decoding neural activity, we simulated a Gaussian random walk with drift, in which each step size was independently drawn from a Gaussian distribution with mean equal to the actual speed and standard deviation equal to the average uncertainty in speed estimates.

## Acknowledgements

We thank Jing Lin and Jian Chen for assistance in stimulus programming. This work was supported by the Simons Collaboration on the Global Brain #324143 and NIH DC004260.

## Author contributions

EA, KJL, GCD and DEA designed the experiment. EA and KJL conducted the experiment and analyzed the data. EA, KJL GCD and DEA wrote the manuscript.

---

**Figure S1.**
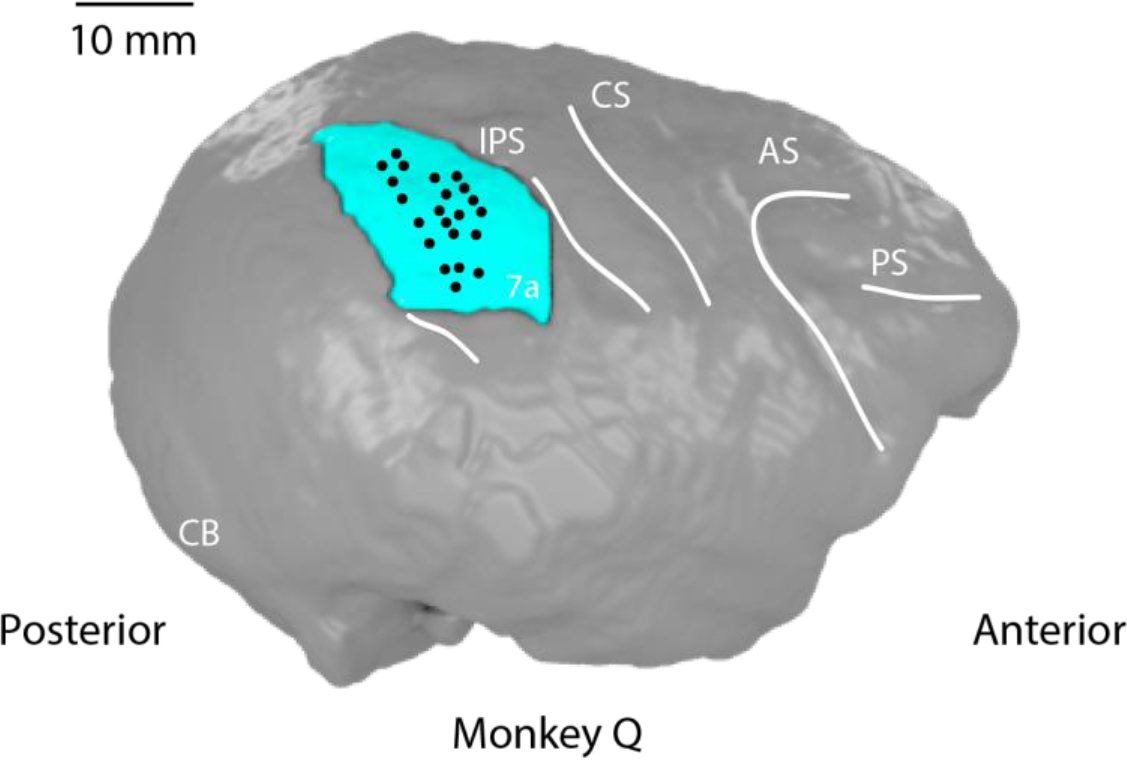
Recording site reconstruction. Magnetic resonance imaging (MRI) reconstruction of recording sites for monkey Q using a frameless stereotaxic technique from two data sets (**Methods**). Right lateral view with area 7a in cyan. Black dots indicate anatomical sites of recordings in area 7a (22 recording sessions). AS, arcuate sulcus; CB, cerebellum; CS, central sulcus; IPS, intraparietal sulcus; PS, principal sulcus.

**Figure S2.**
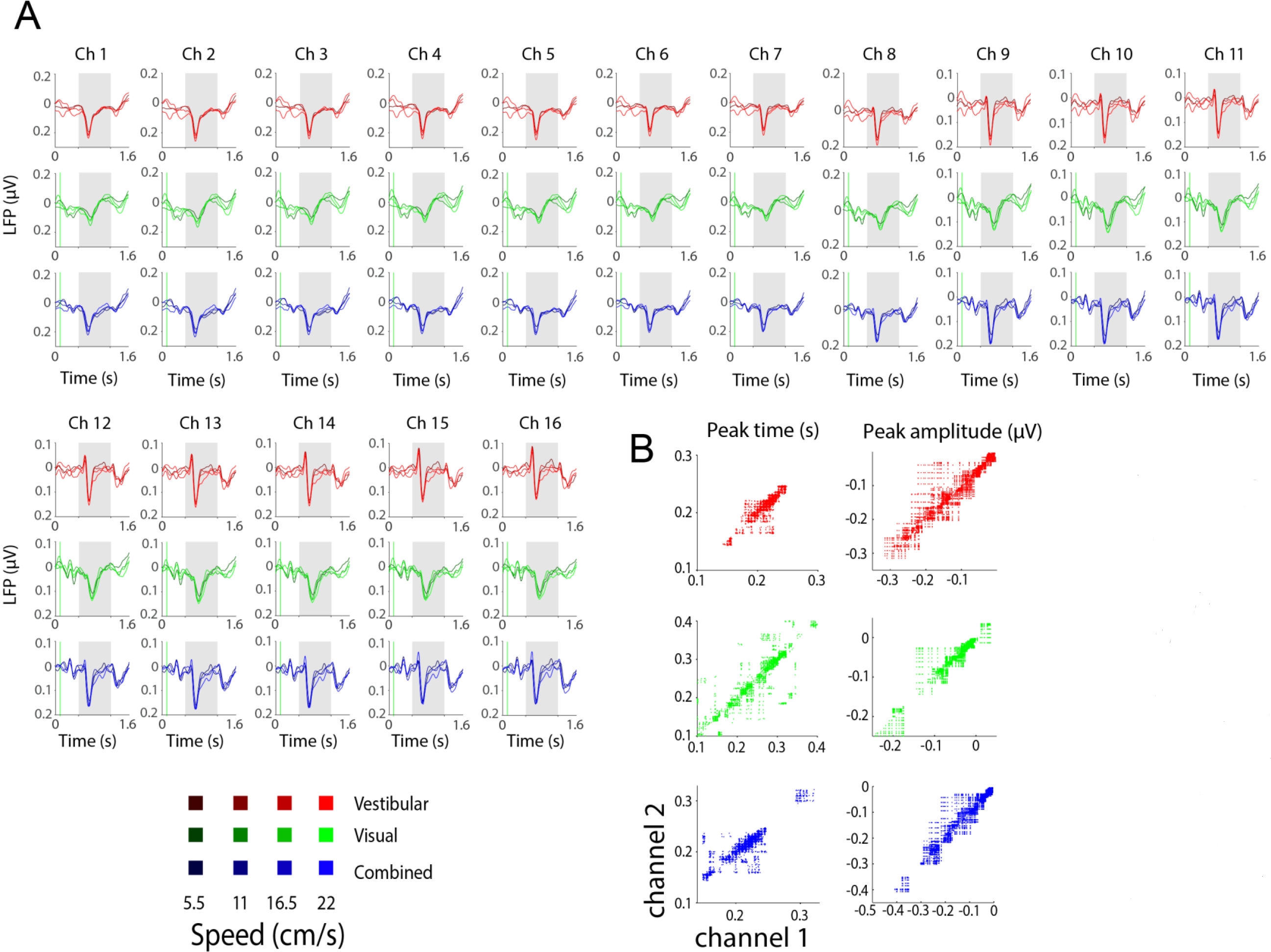
Local-field potentials are spatially correlated. **(A)** Stimulus-evoked local field potential (LFP) recorded from all channels (Ch 1-16) of a linear array during one example recording session. Each panel shows trial-averaged LFP traces in response to four different speeds of linear translation for vestibular (red), visual (green) and combined conditions (blue). Gray shaded regions correspond to periods of selfmotion. **(B)** Comparison of the timing (*left*) and amplitude (*right*) of the dominant peaks or troughs of the stimulus-evoked potentials for all simultaneously recorded pairs of electrodes across all recording sessions (*n*=5280). Both measures are highly correlated across recording sites.

**Figure S3.**
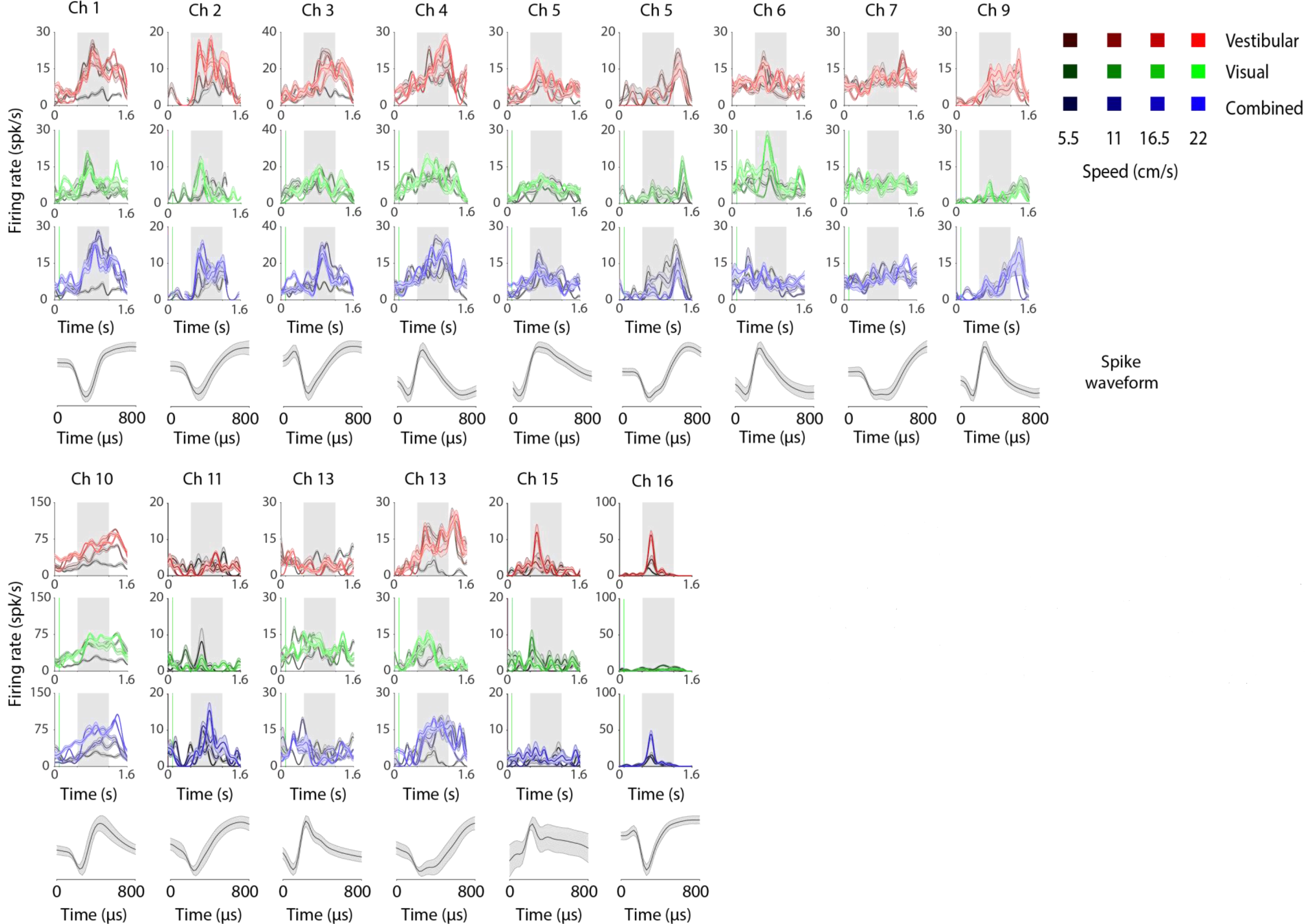
Diversity of neuronal responses to linear translation. Average time course of responses of fifteen single-units isolated during the same recording session as in **Figure S2**. Shaded error bars denote ± 1 SEM. The bottom-most panel of each column shows the average waveform (mean ± 1 standard deviation) from the corresponding single-unit. Note that not all units respond to all speeds and conditions, and not all units have a sustained activity during the motion period.

**Figure S4.**
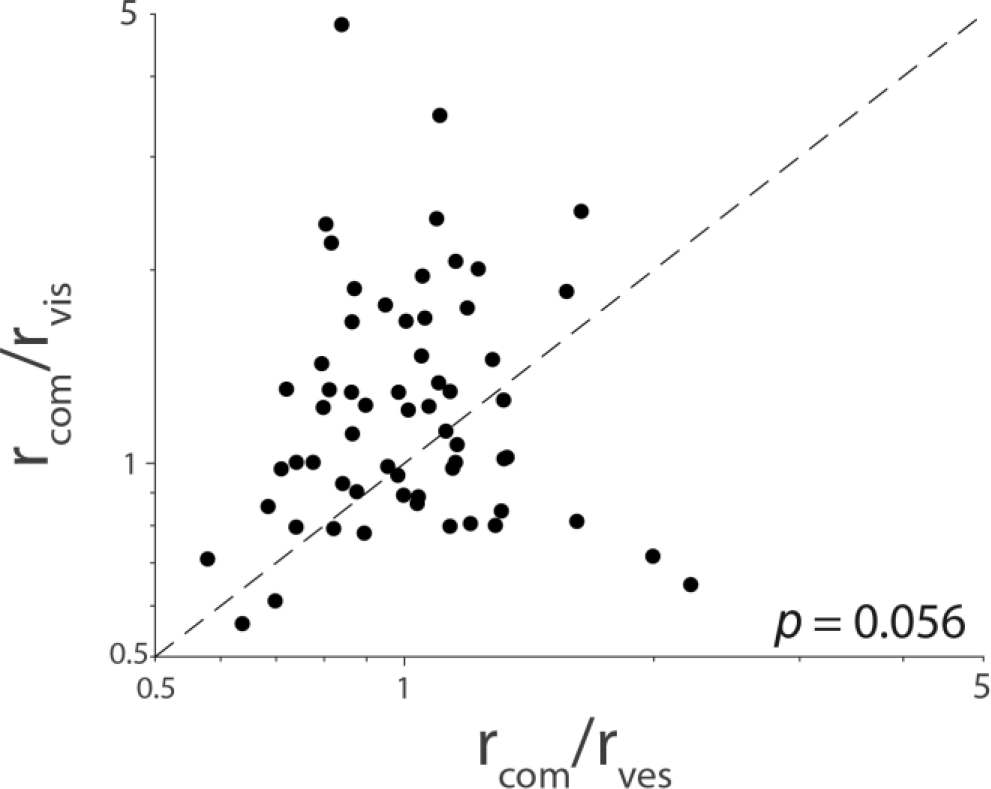
Modulation of firing rates by sensory modality. Comparison of the ratio of peak response elicited by the combined (visual+vestibular) stimulus to peak response elicited by the vestibular (abscissa) or visual (ordinate) motion alone. Data come from the subset of bi-sensory neurons that were responsive to both visual and vestibular inputs (*n*=65). Inset shows p-value (Wilcoxon rank-sum test) for the difference between the median ratios.

**Figure S5.**
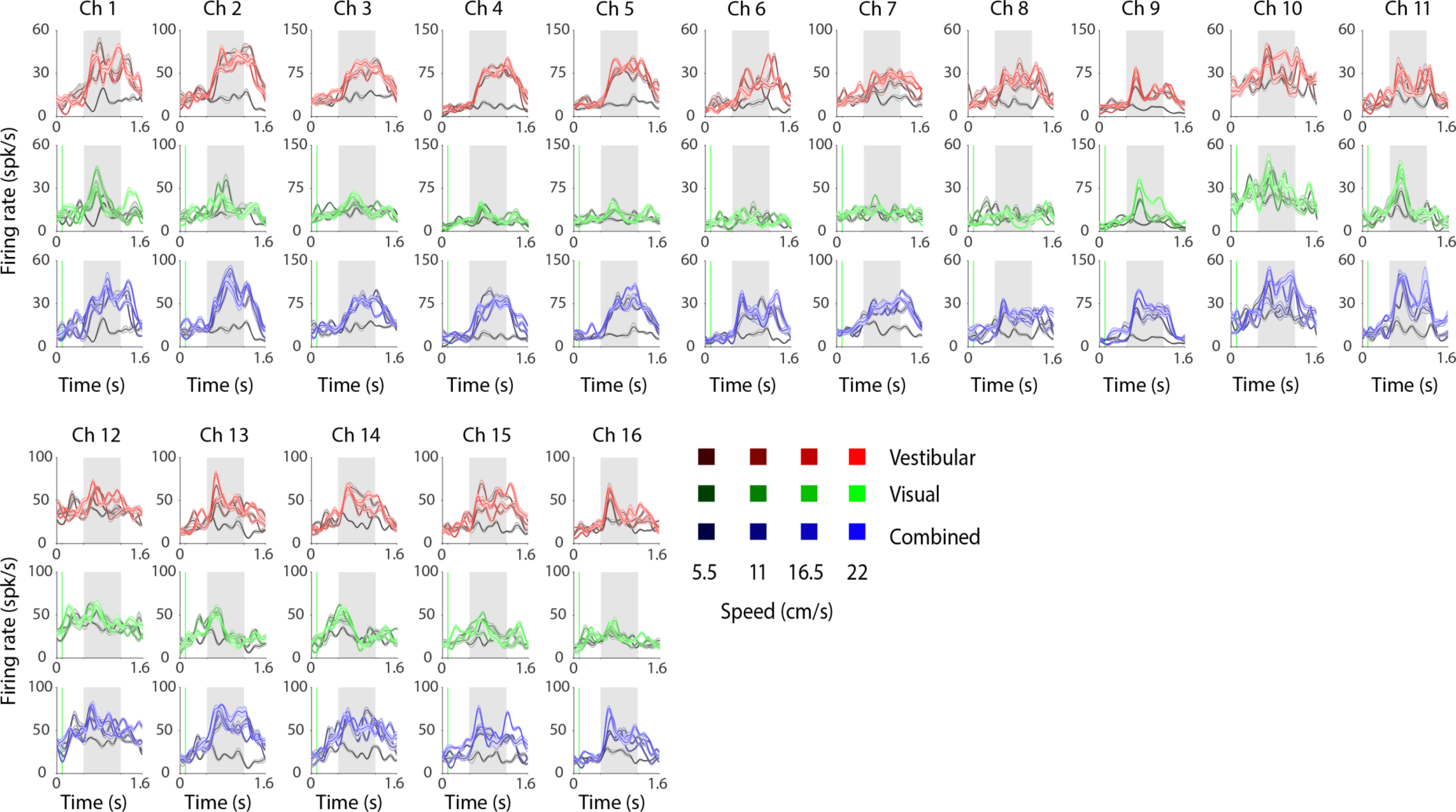
Diversity of multi-unit responses to linear translation. Average time-course of multi-unit responses in all 16 channels during the same recording session as in **Figures S2** and **S3**. Note the similarity of responses elicited by the combined and vestibular stimulus conditions.

**Figure S6.**
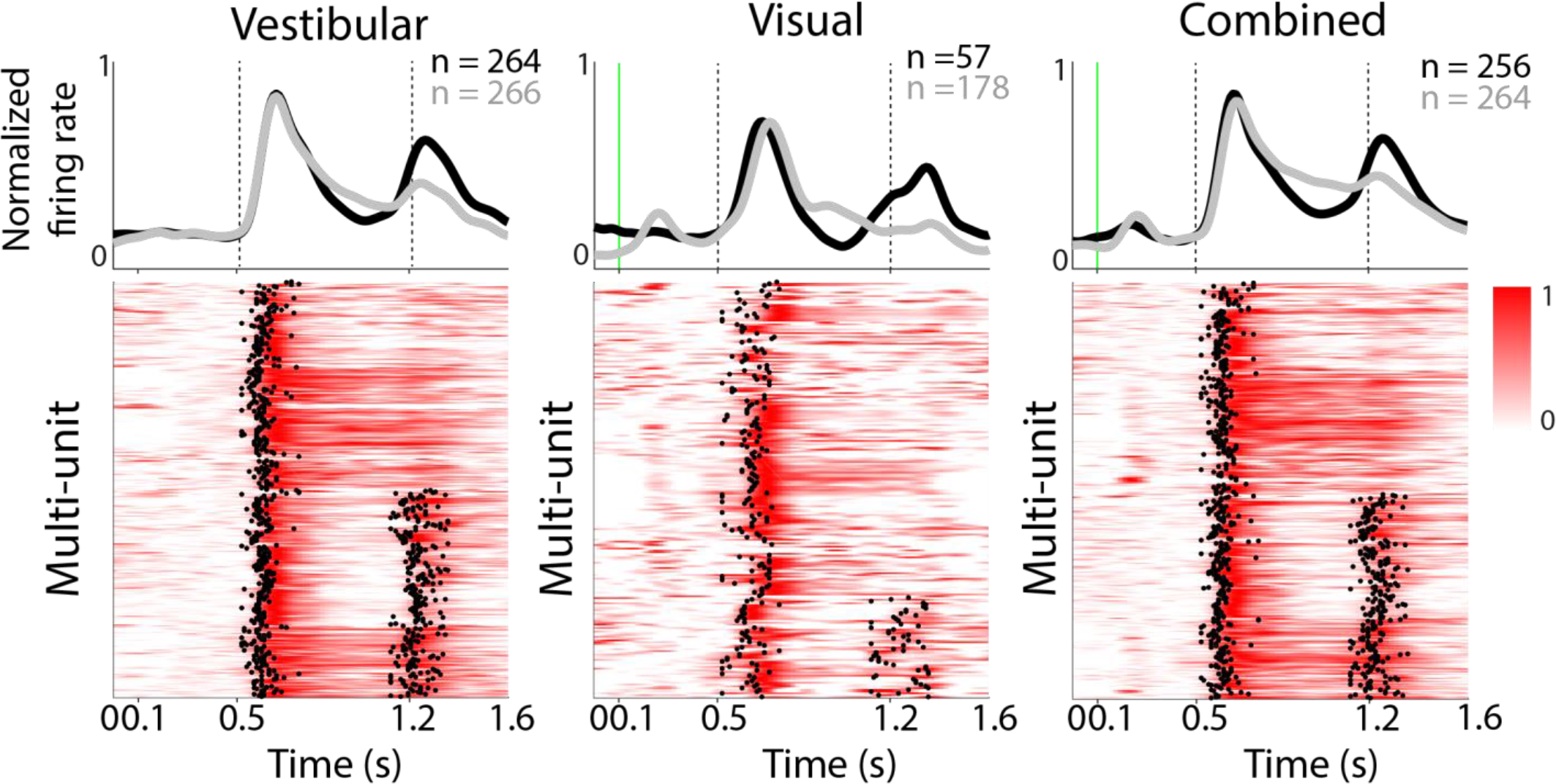
Temporal dynamics of multi-unit responses. **(A)** Each row in the colored panels shows the trial- averaged excitatory response of multi-units (MUA) to vestibular (left, *n*=530), visual (middle, *n*=235) and combined (right, *n*=520) linear translation stimuli. Multi-units with unimodal and bimodal responses were averaged separately and are shown on top of the panels (gray vs. black, respectively). Vertical dotted lines show motion onset and offset. The small firing rate increase before motion onset in the visual and combined conditions is due to the cells’ visual response when the stationary dot background is presented.

**Figure S7.**
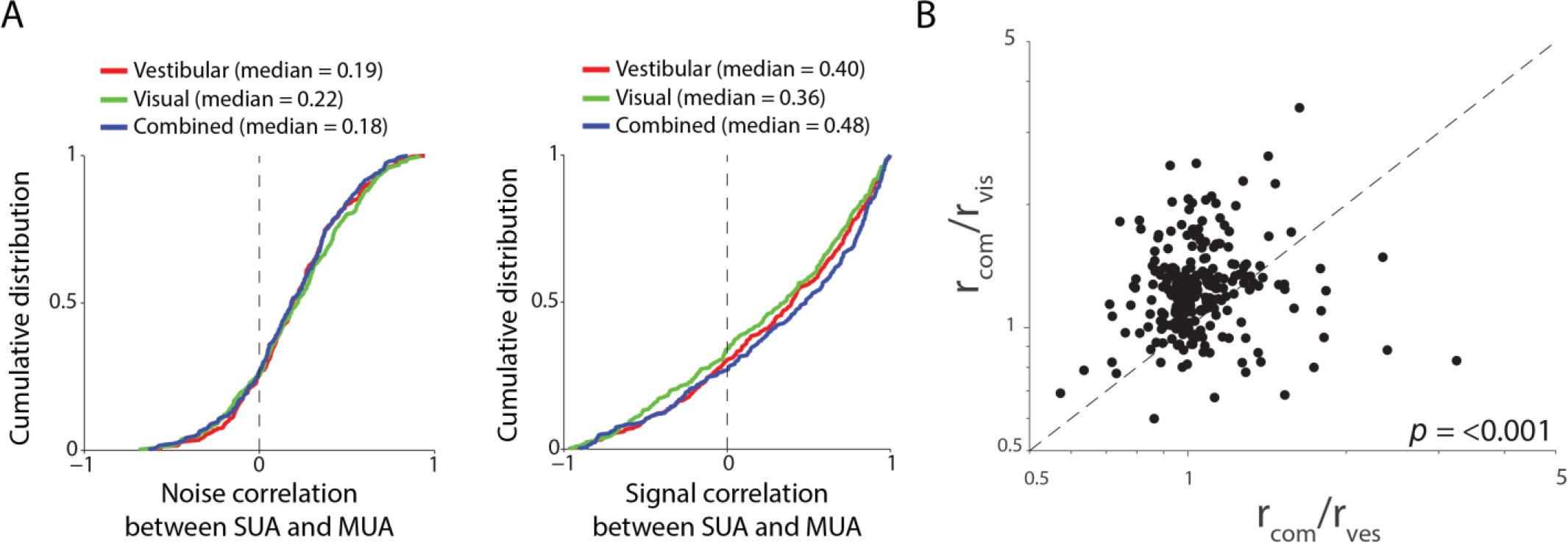
Modulation of multi-unit firing rates by sensory modality. **(A)** Cumulative distribution of correlation coefficients for noise (left) and signal (right) correlations between all pairs of single-unit (SUA) and multi-unit (MUA) responses recorded from the same electrode site. **(B)** Comparison of the ratio of peak response elicited by the combined stimulus to peak response elicited by vestibular (abscissa) or visual (ordinate) motion alone. Data are shown for the subset of bi-sensory multi-units that were responsive to both visual and vestibular inputs (*n*=258). Inset shows p-value (Wilcoxon rank-sum test) for the difference between the median ratios.

**Figure S8.**
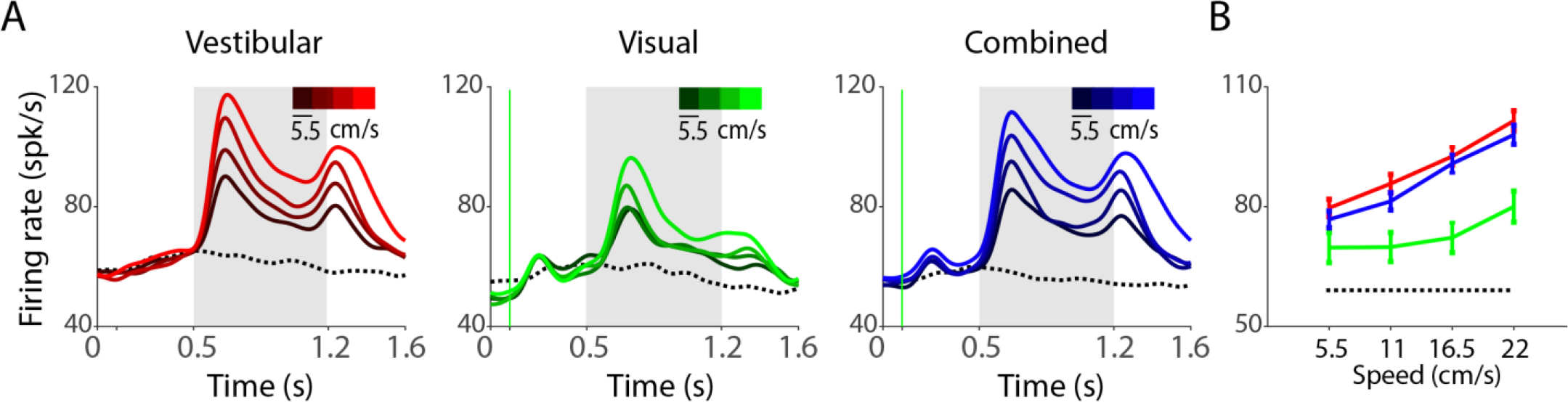
Multi-unit responses increase with translation speed. **(A)** Time course of response averaged across multi-units tuned to translation speed under the vestibular (red, **n*=530*), visual (green, **n*=235*) and combined (blue, **n*=520*) conditions. Different hues correspond to different speeds of motion (dark-slower speeds, brigh*t*-higher speeds). Dotted line denotes response to the baseline condition. Grey shaded areas show periods of motion. **(B)** Population average tuning curves of the multi-units shown in (A).

**Figure S9.**
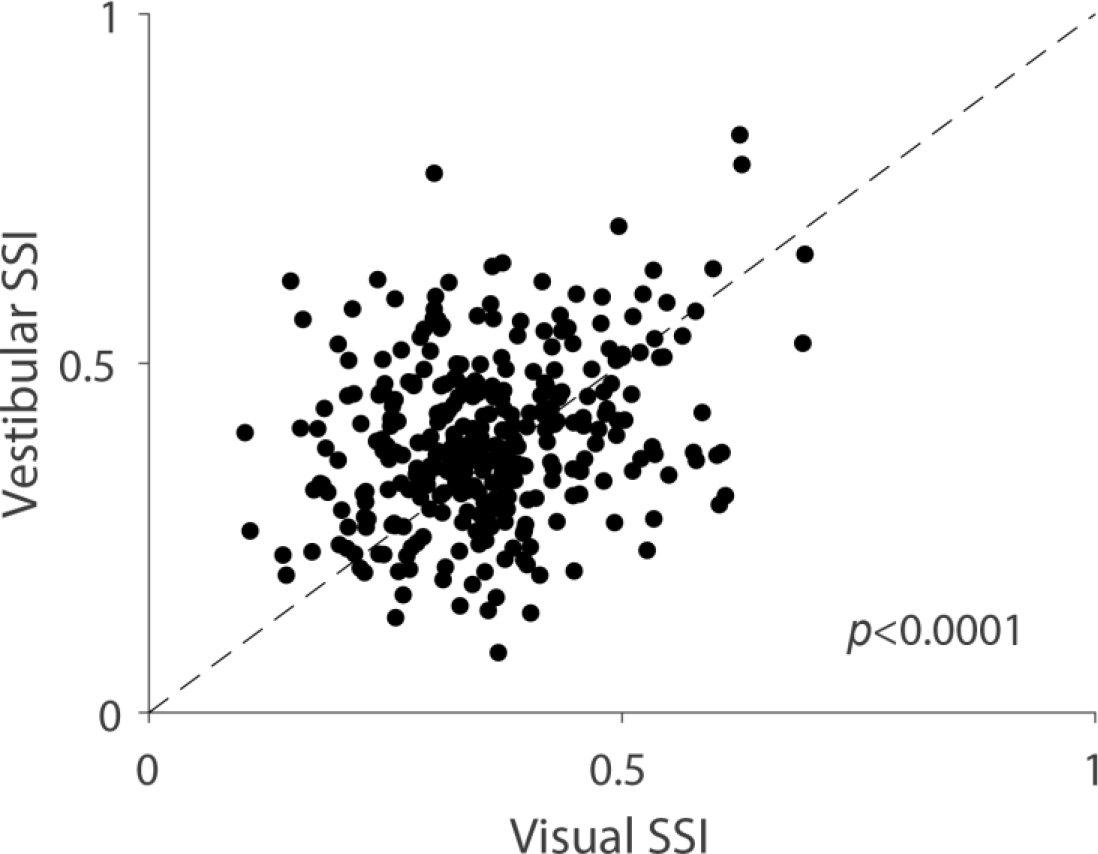
Speed selectivity indices differ across stimulus conditions. Speed selectivity indices (SSI) to linear speed for all isolated single neurons (*n*=340) tested under the vestibular (ordinate) and visual (abscissa) conditions. The median SSI is significantly greater for the vestibular condition (*t*-test, *p*=1.64x10^−5^).

**Figure S10.**
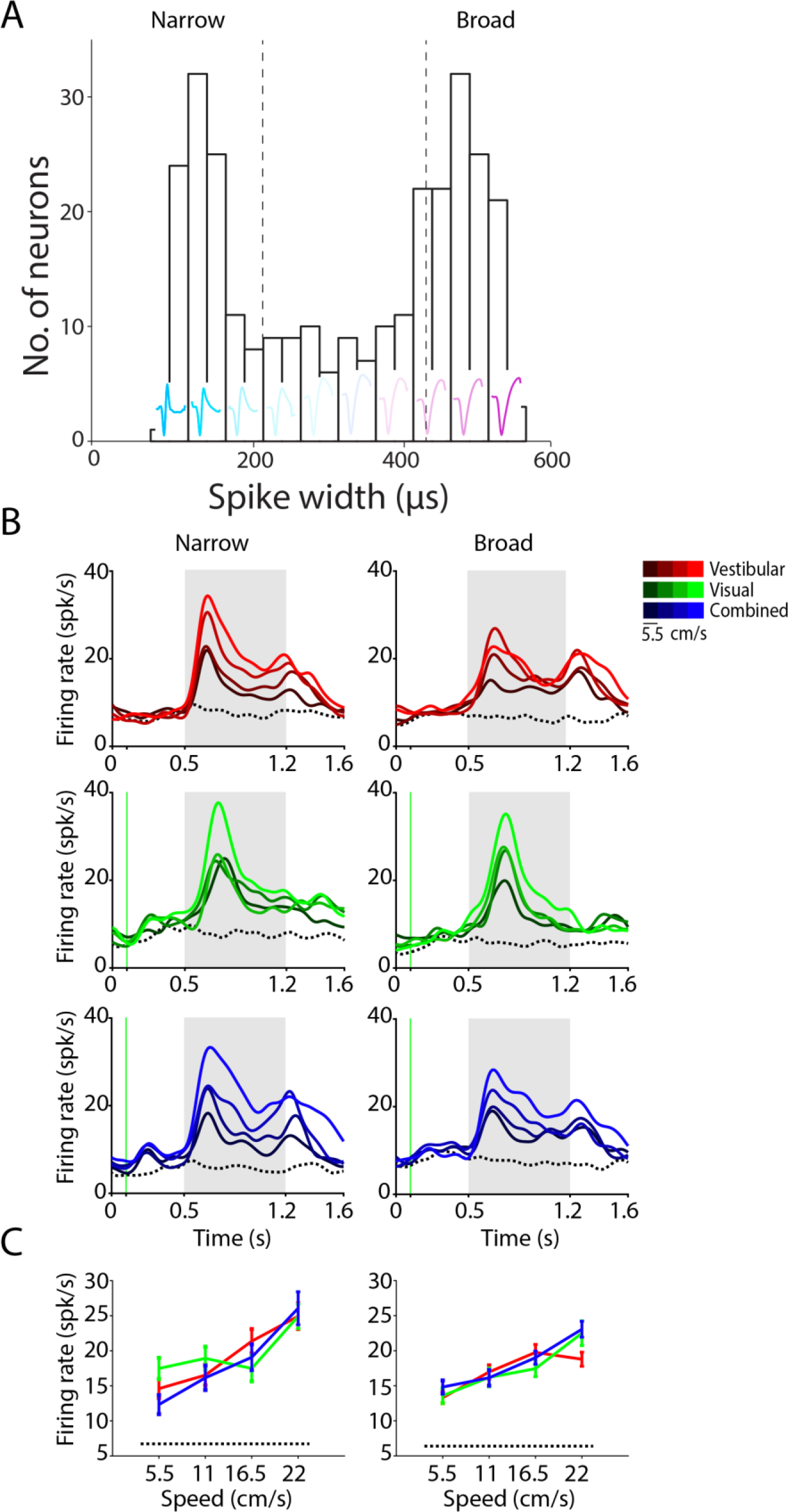
Narrow and broad-spiking neurons have qualitatively similar responses. **(A)** Distribution of spike-widths across the population of all neurons was found to be significantly different from unimodal (Hartigan’s dip-test; *p*=0.036). The average waveforms of all neurons in each bin are overlaid. Dashed lines delineate 33^rd^ and 66^th^ percentiles. **(B)** The time-course of average population responses were largely similar for broad-spiking (> 66^th^ percentile) and narrow-spiking (< 33^rd^ percentile) neurons under vestibular (red, narrow **n*=19*, broad **n*=16*), visual (green, narrow **n*=7*, broad **n*=13*) and combined (blue, narrow **n*=13*, broad **n*=18*) conditions. **(C)** Average speed tuning curves for single-units with narrow (left) and broad (right) spike-widths.

**Figure S11.**
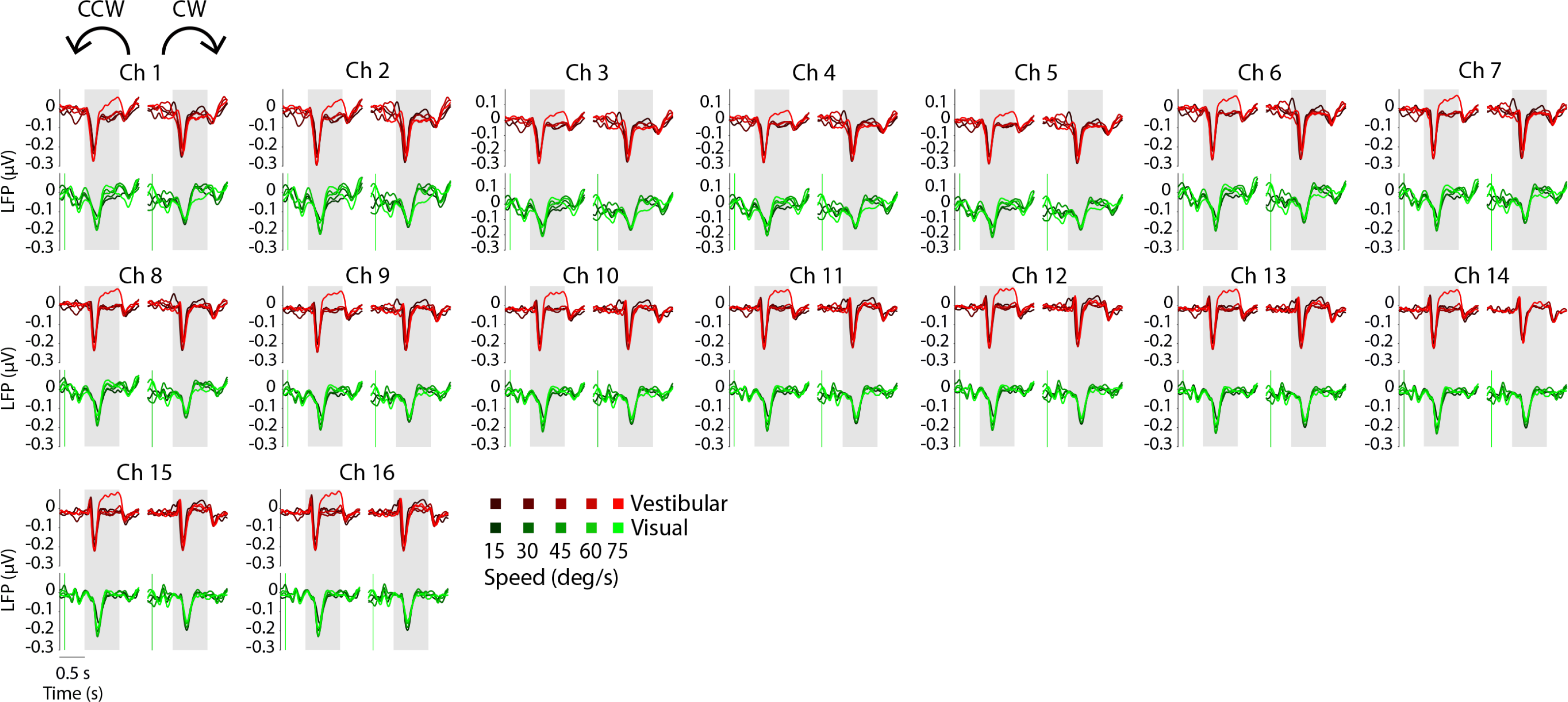
Local field potential (LFP) responses to angular rotation. Stimulus-evoked LFPs recorded simultaneously from the 16 channels of a linear electrode array during angular motion. For each channel, responses to CCW (left) and CW (right) rotations are shown separately for trials with vestibular (red) and visual (green) rotations. Different hues denote different speeds (dark-slower speeds, brigh*t*-faster speeds). Grey shaded areas show periods of motion.

**Figure S12.**
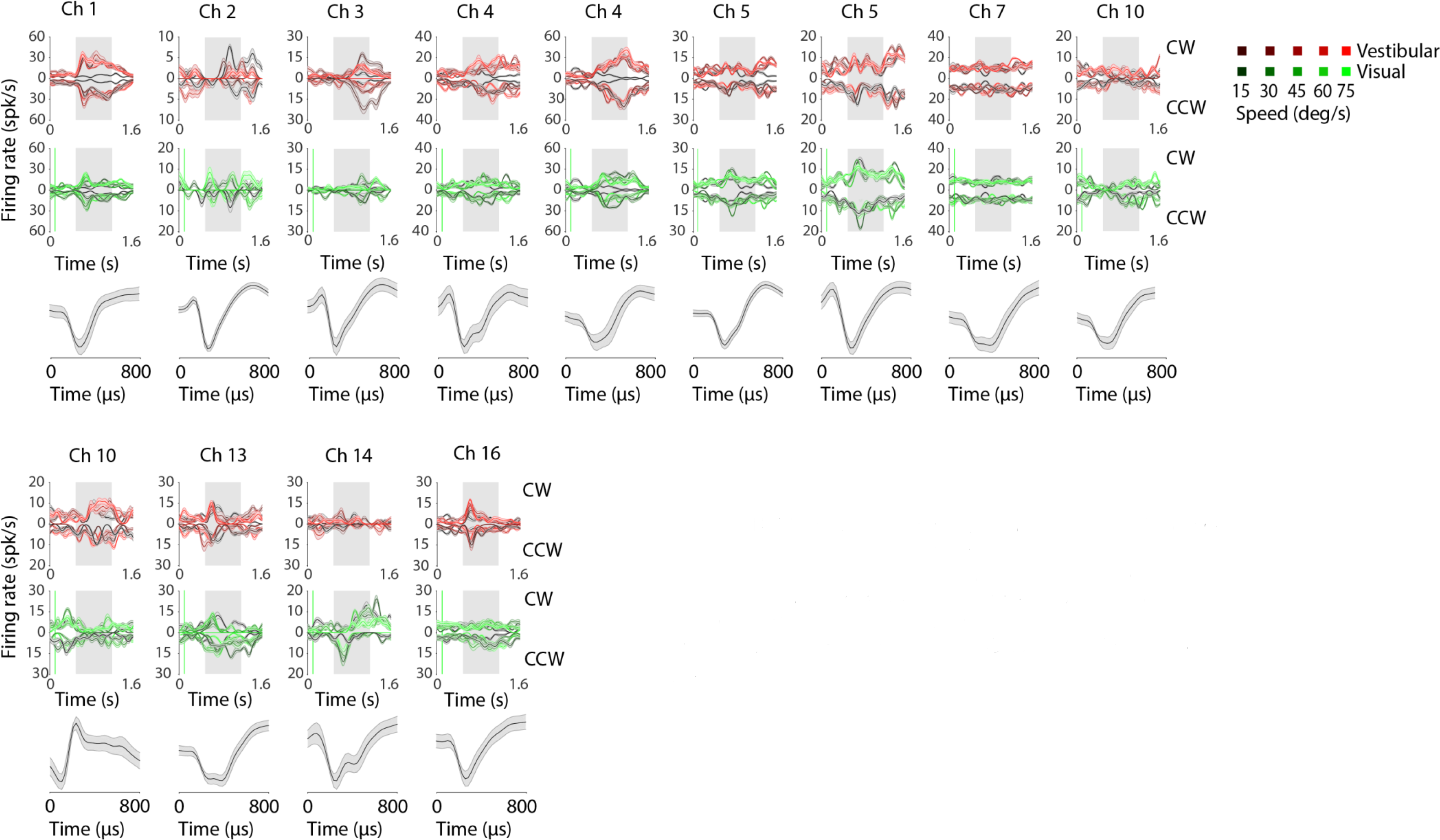
Neuronal responses to angular rotation. Response of the set of all simultaneously recorded single neurons during the same recording session as **Figure S11**. For each channel, responses to rotations are averaged separately and plotted CW (upright axis) and CCW (inverted axis). Different shades correspond to different speeds. Gray shaded areas show time- periods of rotation.

**Figure S13.**
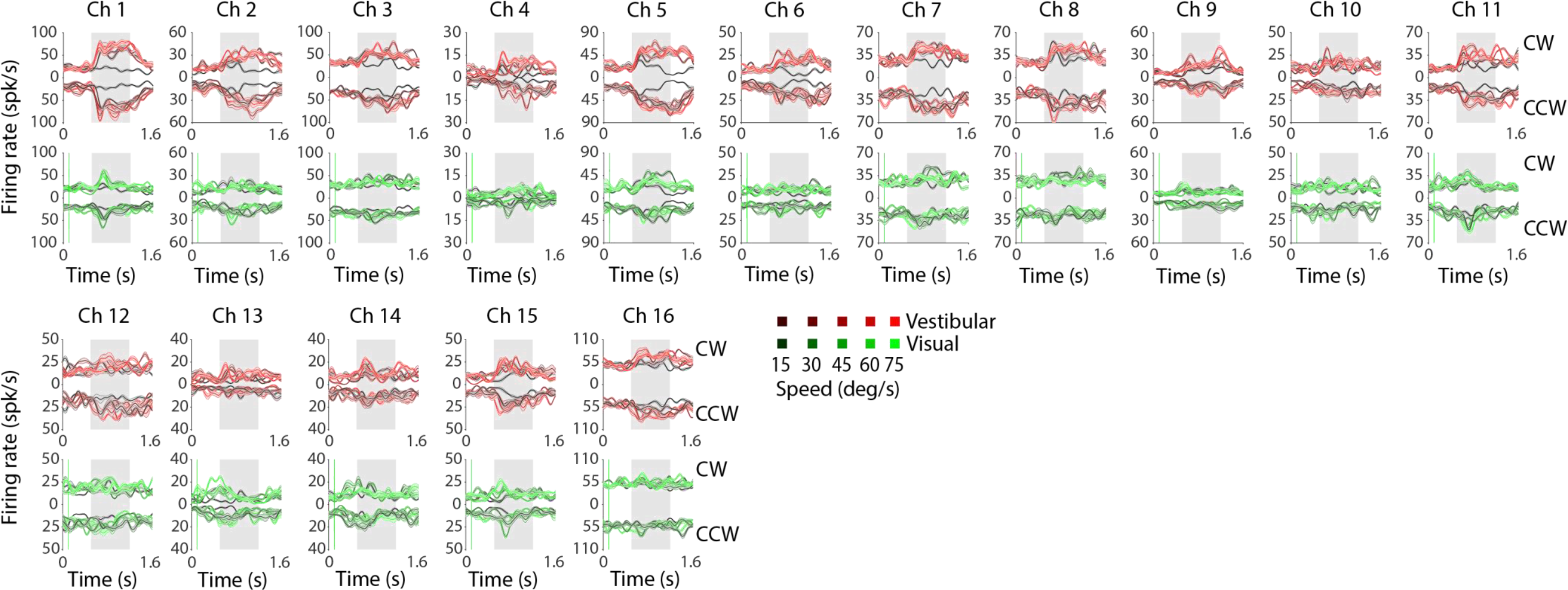
Multi-unit responses to angular rotation. Response of multi-units recorded from all channels during the same recording session as **Figure S11** and **S12**. Conventions are the same as used for plotting single-unit responses in **Figure S12**.

**Figure S14.**
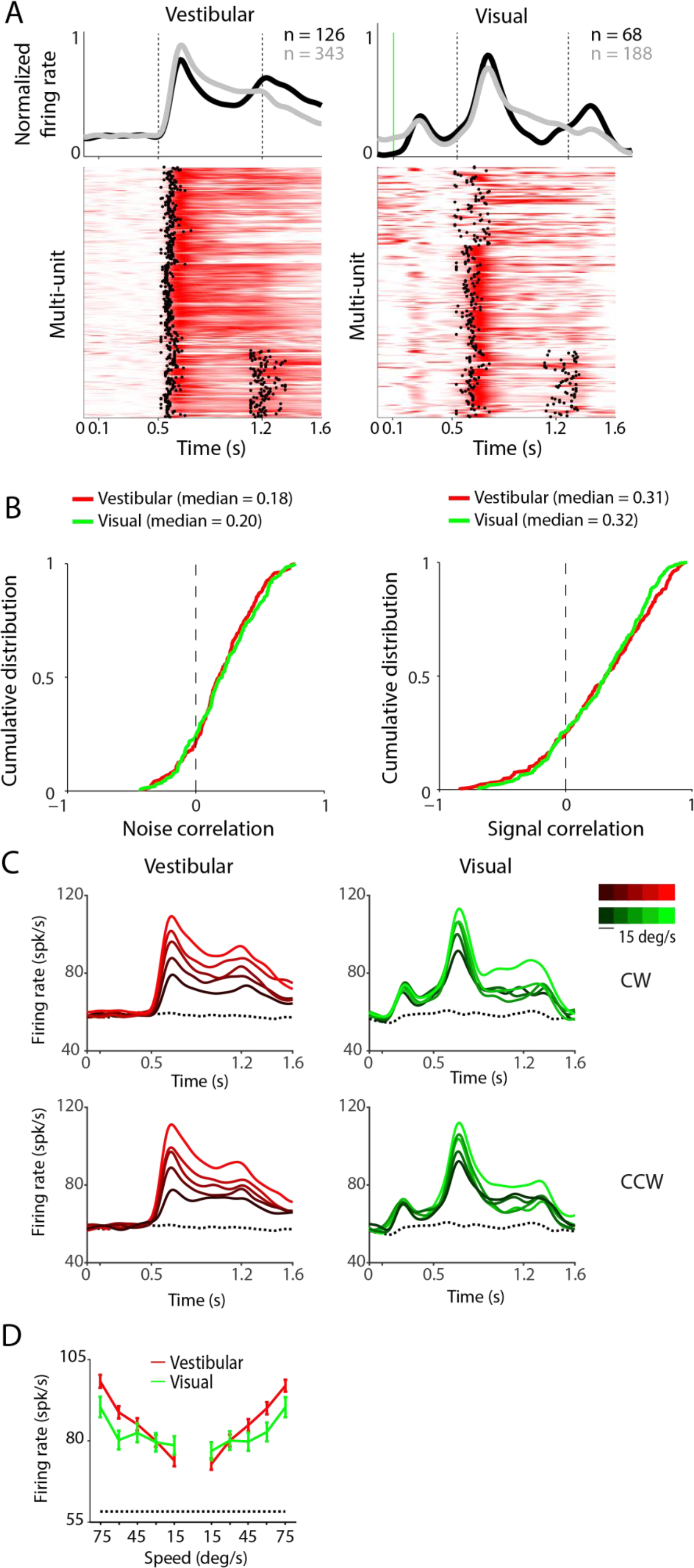
Temporal dynamics and speed selectivity of multi-unit responses to angular rotation. **(A)** Each row in the colored panels shows the trial-averaged response of a multiunit to vestibular (*left*) and visual (*right*) rotation cues. Data from multi-units with unimodal (gray) and bimodal responses (black) were averaged separately and are shown on top of the panels. **(B)** Cumulative distributions of correlation coefficients computed during the motion stimulus, for all pairs of single-unit and multi-unit responses. **(C)** Time course of response averaged across all responsive multi-units with excitatory responses to vestibular (red, **n*=264*) or visual (green, **n*=88*) rotation. Different hues correspond to different speeds of motion (dark-slower speeds, brigh*t*-faster speeds). Dotted line denotes response to the baseline condition. Grey shaded areas show periods of motion. **(D)** Population average tuning curves of the neurons shown in (A) and (B).

**Figure S15.**
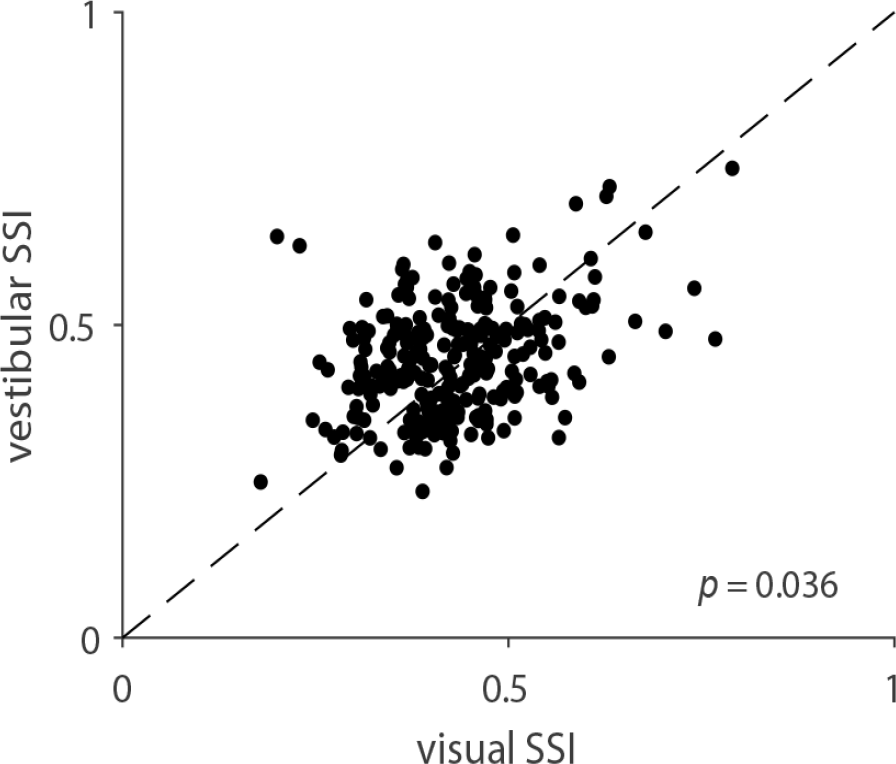
Comparison of speed selectivity indices (SSI) of single-unit responses to vestibular and visual rotation stimuli. SSI values are shown for all isolated single-units (*n*=268).

**Figure S16.**
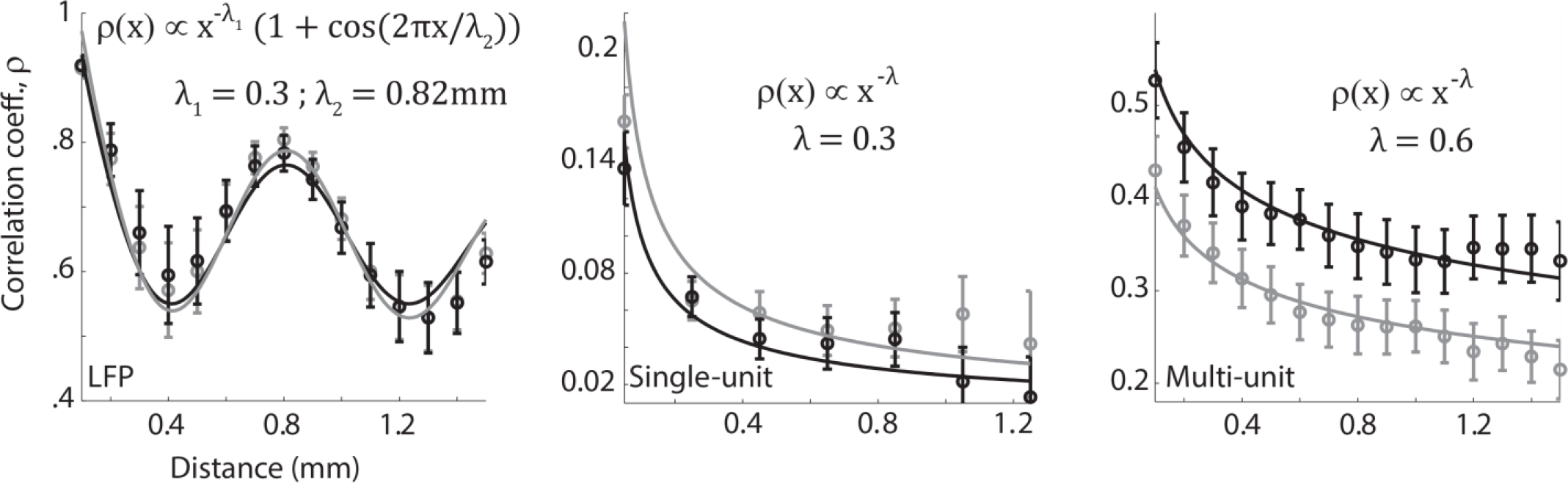
Power-law model explains the spatial dependence of temporal response correlations. Dependence of the pairwise correlation coefficients of LFPs (*left*), single-units (*middle*), and multi-units (*right*) on distance between recording sites was fit to a power-law function. Solid lines correspond to the bes*t*-fit functions to data from linear translation (gray) and angular rotation (black). The functions were fit by constraining the exponent to be identical for both datasets (translation and rotation), but allowing different constants of proportionality. To capture the spatial periodicity of correlations in the LFP, we added a sinusoidal wave function with wavelength being the only other free parameter (phase was set to zero). Model parameters are shown on top of each panel correspond to the best fit values across both protocols.

**Figure S17.**
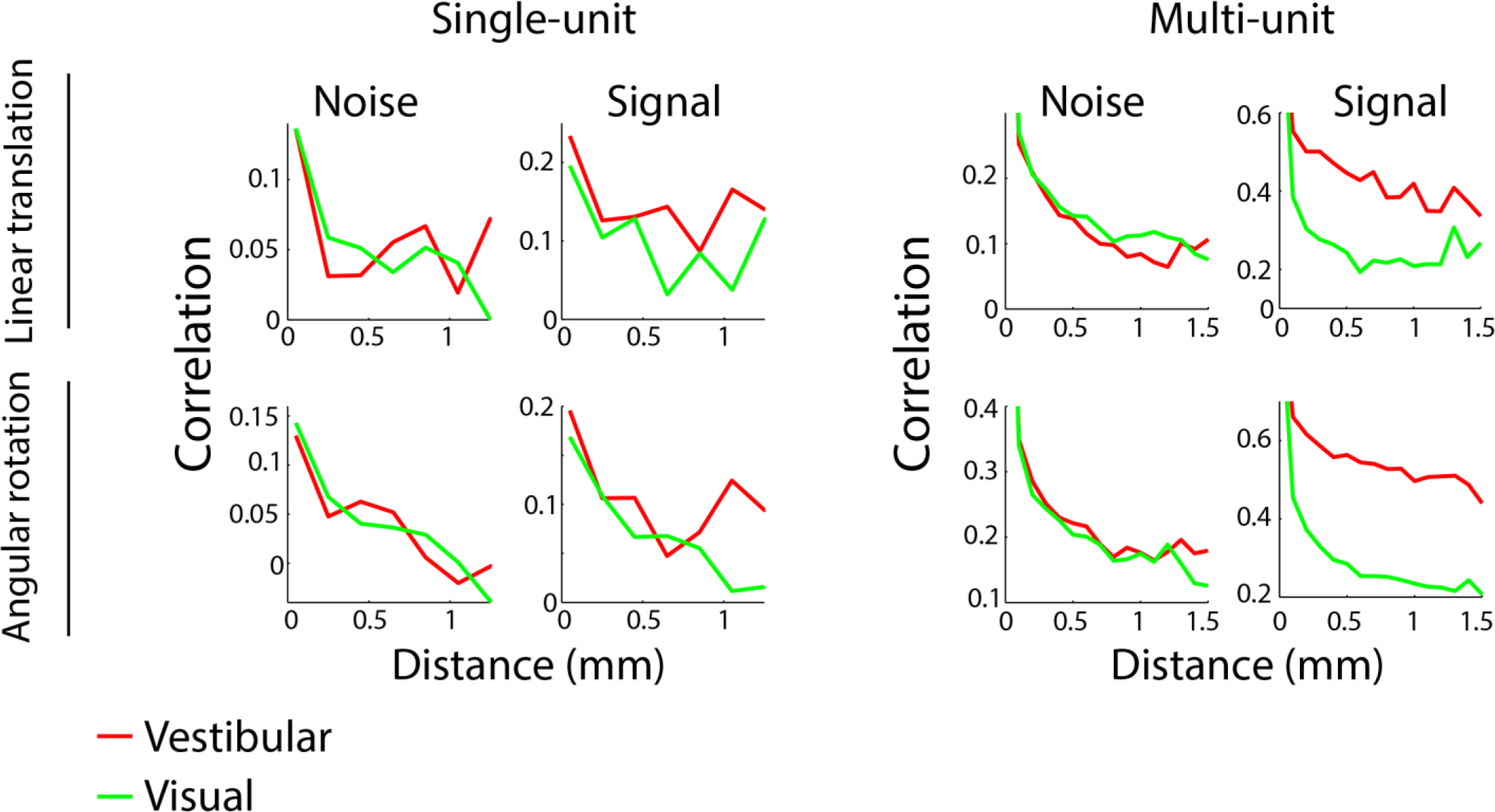
Spatial profile of correlated variability in spike counts. Correlations in spike-count variability between pairs of single-units (left) or pairs of multi-units (right) induced by the stimulus (signal correlation) or other factors (noise correlation) for all simultaneously recorded pairs in our data set (single-units: *n*=1537 pairs, multi-units: *n*=5280 pairs). The top row shows data for the linear translation condition, whereas the bottom row shows data for angular rotation.

**Figure S18.**
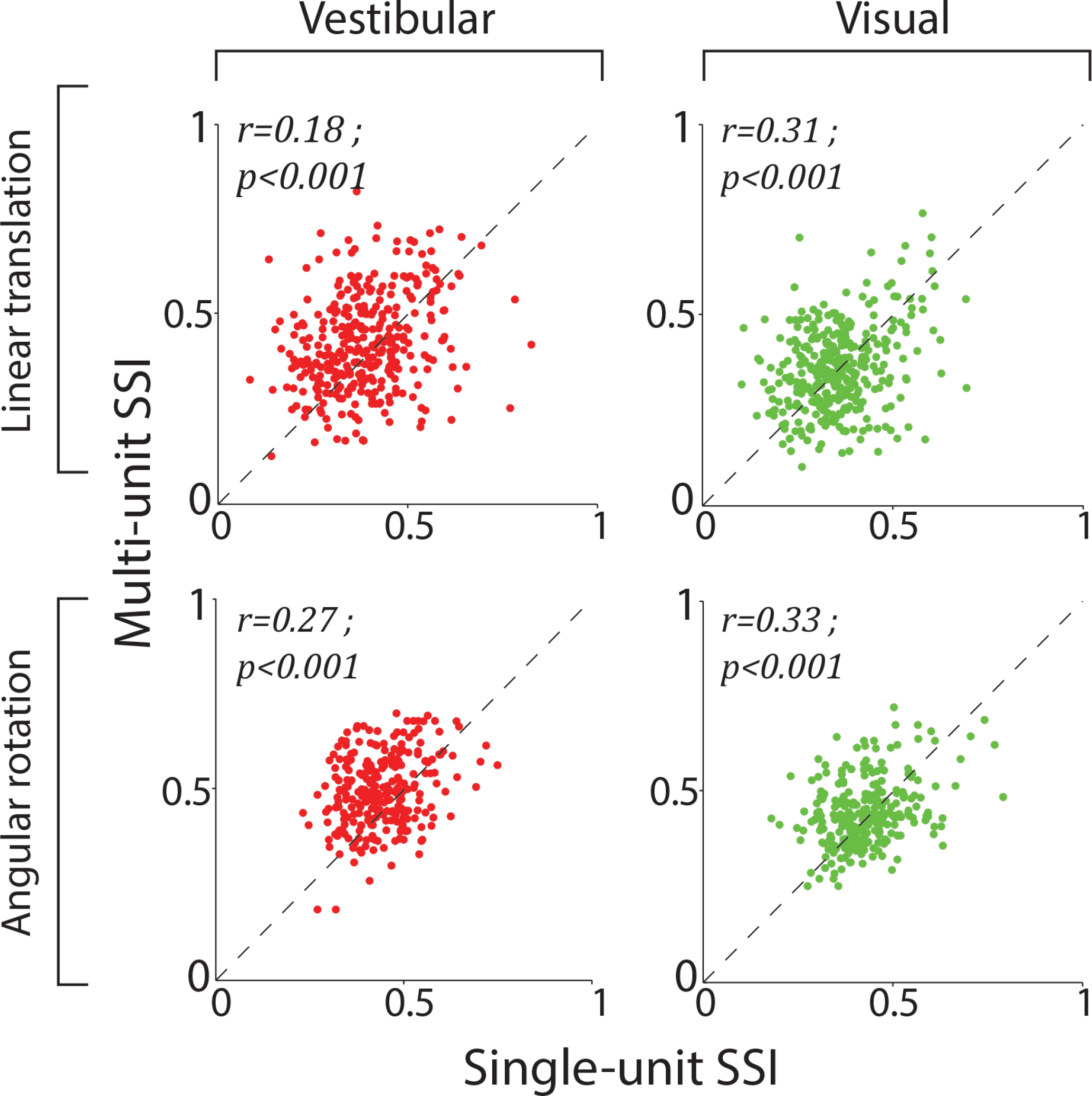
Comparison of single- and multi-unit speed selectivity indices (SSIs). Comparison of SSI values computed from single- and multi-unit activity for all pairs of signals recorded from the same channel. Data are shown for translation (top, *n*=340) and rotation (bottom, *n*=268) stimuli under the vestibular (left) and visual (right) conditions. Pearson’s correlation coefficients (*r*) and its p-value are shown on top of each panel.

**Figure S19.**
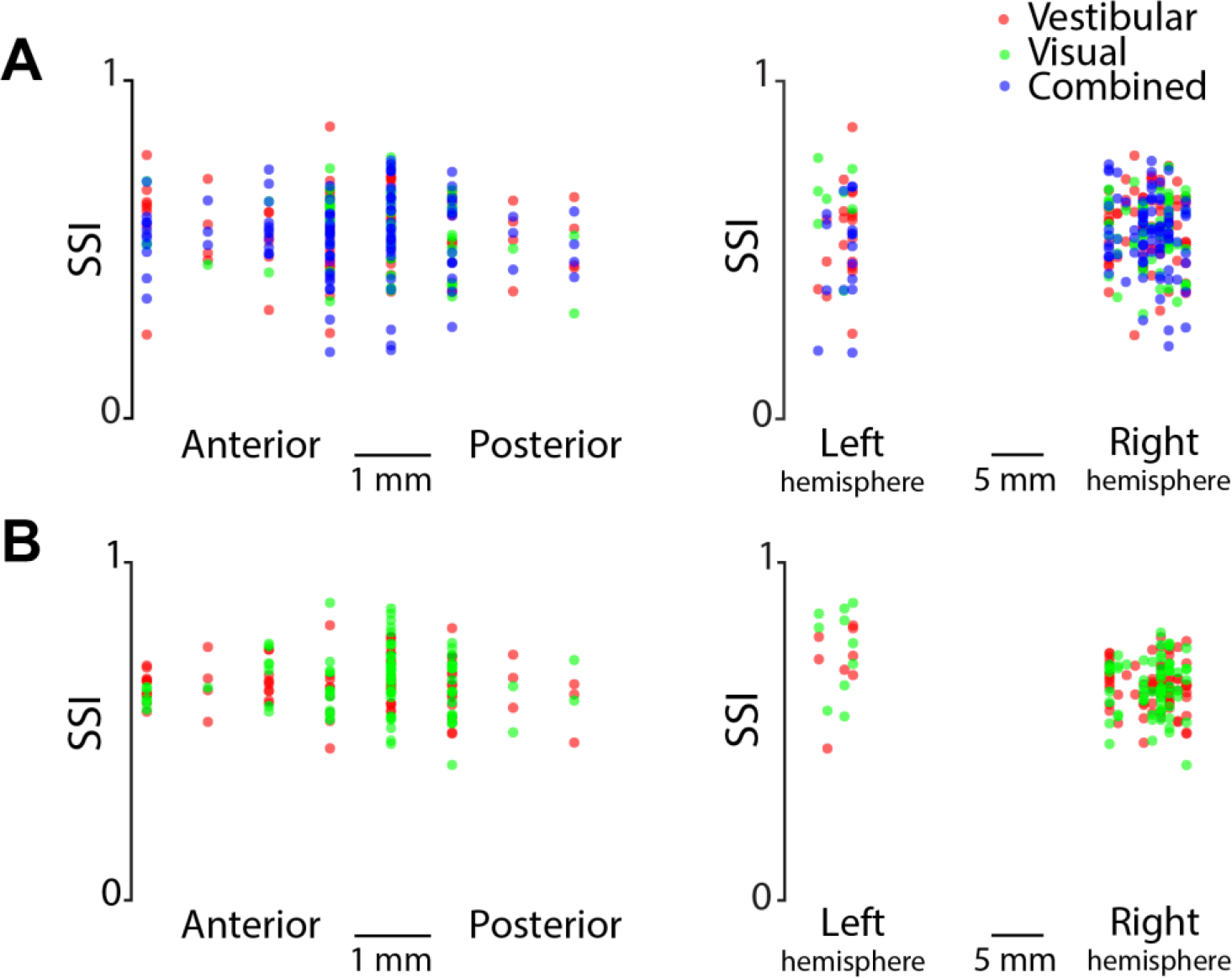
Strength of speed tuning does not depend on recording location. **(A)** Speed selectivity indices (SSI) for all neurons responsive to linear translation, plotted as a function of the relative coordinates of recording sites along the anterior/posterior axis (*left*) and lateral/medial (*right*) axes. **(B)** Similar to (A), but for neurons responsive to angular rotation.

## References

Alexander, A. S., & Nitz, D. a. (2015). Retrosplenial cortex maps the conjunction of internal and external spaces. Nature Neuroscience, 18(July), 1–12.

Andersen, R. A. (1989). Visual and eye movement functions of the posterior parietal cortex. Annual Review of Neuroscience, 12(1), 377–403.

Andersen, R. A., Asanuma, C., Essick, G., & Siegel, R. M. (1990). Corticocortical connections of anatomically and physiologically defined subdivisions within the inferior parietal lobule. Journal of Comparative Neurology,296(1), 65–113.

Andersen, R. A., Essick, G. K., & Siegel, R. M. (1985). Encoding of spatial location by posterior parietal neurons. Science, 230(4724), 456–8.

Barrow, C. J., & Latto, R. (1996). The role of inferior parietal cortex and fornix in route following and topographic orientation in cynomolgus monkeys. Behavioural Brain Research, 75(1–2), 99–112.

Berens, P., Ecker, A. S., Cotton, R. J., Ma, W. J., Bethge, M., & Tolias, A. S. (2012). A Fast and Simple Population Code for Orientation in Primate V1. Journal of Neuroscience, 32(31), 10618–10626.

Bremmer, F., Duhamel, J. R., Ben Hamed, S., & Graf, W. (2002). Heading encoding in the macaque ventral intraparietal area (VIP). European Journal of Neuroscience, 16(8), 1554–1568.

Bremmer, F., Klam, F., Duhamel, J.-R., Ben Hamed, S., & Graf, W. (2002). Visual-vestibular interactive responses in the macaque ventral intraparietal area (VIP). The European Journal of Neuroscience, 16(8), 1569–86.

Britten, K. H. (2008). Mechanisms of Self-Motion Perception. Annual Review of Neuroscience, 31(1), 389–410.

Britten, K. H., & van Wezel, R. J. A. (1998). Electrical microstimulation of cortical area MST biases heading perception in monkeys. Nature Neuroscience, 1(1), 59–63.

Britten, K. H., & Van Wezel, R. J. A. (2002). Area MST and heading perception in macaque monkeys. Cerebral Cortex, 12(7), 692–701.

Brotchie, P. R., Andersen, R. A., Snyder, L. H., & Goodman, S. J. (1995). Head position signals used by parietal neurons to encode locations of visual stimuli. Nature, 375(6528), 232–235.

Bucci, D. J. (2009). Posterior parietal cortex: An interface between attention and learning? Neurobiology of Learning and Memory, 91(2), 114–120.

Burgess, N. (Neil), Jeffery, K. J., O’Keefe, J. (1999). The hippocampal and parietal foundations of spatial cognition. Oxford University Press. Great Britain

Cavada, C., & Goldman-Rakic, P. S. (1989). Posterior parietal cortex in rhesus monkey: II. Evidence for segregated corticocortical networks linking sensory and limbic areas with the frontal lobe. Journal of Comparative Neurology, 287(4), 422–445.

Chafee, M. V, & Crowe, D. a. (2012). Thinking in spatial terms: decoupling spatial representation from sensorimotor control in monkey posterior parietal areas 7a and LIP. Frontiers in Integrative Neuroscience, 6(January), 112.

Chen, A., DeAngelis, G. C., & Angelaki, D. E. (2010). Macaque Parieto-Insular Vestibular Cortex: Responses to Self-Motion and Optic Flow. Journal of Neuroscience, 30(8), 3022–3042.

Chen, A., DeAngelis, G. C., & Angelaki, D. E. (2011). A comparison of vestibular spatiotemporal tuning in macaque parietoinsular vestibular cortex, ventral intraparietal area, and medial superior temporal area. The Journal of Neuroscience, 31(8), 3082–3094.

Chen, A., DeAngelis, G. C., & Angelaki, D. E. (2013). Functional Specializations of the Ventral Intraparietal Area for Multisensory Heading Discrimination. Journal of Neuroscience, 33(8), 3567–3581.

Chen, A., Gu, Y., Takahashi, K., Angelaki, D. E., & Deangelis, G. C. (2008). Clustering of self-motion selectivity and visual response properties in macaque area MSTd. Journal of Neurophysiology, 100(5), 2669–2683.

Chen, L. L., Lin, L. H., Barnes, C. A., & McNaughton, B. L. (1994). Head-direction cells in the rat posterior cortex-II. Contributions of visual and ideothetic information to the directional firing. Experimental Brain Research, 101(1), 24–34.

Chen, L. L., Lin, L. H., Green, E. J., Barnes, C. A., & McNaughton, B. L. (1994). Head-direction cells in the rat posterior cortex-I. anatomical distribution and behavioral modulation. Experimental Brain Research, 101(1), 8–23.

Ciaramelli, E., Rosenbaum, R. S., Solcz, S., Levine, B., & Moscovitch, M. (2010). Mental space travel: damage to posterior parietal cortex prevents egocentric navigation and reexperiencing of remote spatial memories. Journal of Experimental Psychology. Learning, Memory, and Cognition, 36(3), 619–634.

Cohen, Y. E., & Andersen, R. a. (2002). A common reference frame for movement plans in the posterior parietal cortex. Nature Reviews. Neuroscience, 3(7), 553–62.

Constantinidis, C., & Steinmetz, M. A. (1996). Neuronal activity in posterior parietal area 7a during the delay periods of a spatial memory task. Journal of Neurophysiology, 76(2), 1352–1355.

Constantinidis, C., & Steinmetz, M. A. (2001a). Neuronal responses in area 7a to multiple-stimulus displays: I. neurons encode the location of the salient stimulus. Cerebral Cortex (New York, N.Y. : 1991), 11(7), 581–591.

Constantinidis, C., & Steinmetz, M. A. (2001b). Neuronal responses in area 7a to multiple stimulus displays: ii. responses are suppressed at the cued location. Cereb Cortex, 11(7), 592–597.

Cowen, S. L., & Nitz, D. A. (2014). Repeating Firing Fields of CA1 Neurons Shift Forward in Response to Increasing Angular Velocity. The Journal of Neuroscience, 34(1), 232–241.

Ding, S. L., Van Hoesen, G., & Rockland, K. S. (2000). Inferior parietal lobule projections to the presubiculum and neighboring ventromedial temporal cortical areas. Journal of Comparative Neurology, 425(4), 510–530.

Dubowitz, D. J., & Scadeng, M. (2011). A frameless stereotaxic MRI technique for macaque neuroscience studies. The Open Neuroimaging Journal, 5(Suppl 2), 198–205.

Duffy, C. J., & Wurtz, R. H. (1997). Medial superior temporal area neurons respond to speed patterns in optic flow. The Journal of Neuroscience 17(8), 2839–2851.

Einevoll, G. T., Kayser, C., Logothetis, N. K., & Panzeri, S. (2013). Modelling and analysis of local field potentials for studying the function of cortical circuits. Nat Rev Neurosci, 14(11), 770–785.

Etienne, A. S., & Jeffery, K. J. (2004). Path integration in mammals. Hippocampus, 14(2), 180–192.

Etienne, A. S., Maurer, R., & Séguinot, V. (1996). Path integration in mammals and its interaction with visual landmarks. The Journal of Experimental Biology, 199(Pt 1), 201–9.

Faugier-Grimaud, S., & Ventre, J. (1989). Anatomic connections of inferior parietal cortex (area 7) with subcortical structures related to vestibulo-ocular function in a monkey (macaca fascicularis). Journal of Comparative Neurology, 280(1), 1–14.

Freedman, D. J., & Assad, J. A. (2006). Experience-dependent representation of visual categories in parietal cortex. Nature, 443(7107), 85–88.

Frey, S., Pandya, D. N., Chakravarty, M. M., Bailey, L., Petrides, M., & Collins, D. L. (2011). An MRI based average macaque monkey stereotaxic atlas and space (MNI monkey space). NeuroImage, 55(4), 1435–1442.

G. Paxinos, X. F. Huang, A. W. T. (2000). The Rhesus Monkey Brain in Stereotaxic Coordinates. Academic Press. USA.

Glasauer, S., Amorim, M. A., Vitte, E., & Berthoz, A. (1994). Goal-directed linear locomotion in normal and labyrinthine-defective subjects. Experimental Brain Research, 98(2), 323–35.

Graf, A. B. A., Kohn, A., Jazayeri, M., & Movshon, J. A. (2011). Decoding the activity of neuronal populations in macaque primary visual cortex. Nature Neuroscience, 14(2), 239–245.

Gu, Y., Angelaki, D. E., & DeAngelis, G. C. (2008). Neural correlates of multisensory cue integration in macaque MSTd. Nature Neuroscience, 11(10), 1201–1210.

Gu, Y., Watkins, P. V, Angelaki, D. E., & DeAngelis, G. C. (2006). Visual and nonvisual contributions to three-dimensional heading selectivity in the medial superior temporal area. Journal of Neuroscience, 26(1), 73–85.

Hartigan, J. A., & Hartigan, P. M. (1985). The Dip Test of Unimodality. The Annals of Statistics, 13(1), 70–84.

Hinman, J. R., Brandon, M. P., Climer, J. R., Chapman, G. W., & Hasselmo, M. E. (2016). Multiple Running Speed Signals in Medial Entorhinal Cortex. Neuron, 91(3), 666–679.

Israël, I., Grasso, R., Georges-Francois, P., Tsuzuku, T., & Berthoz, A. (1997). Spatial memory and path integration studied by self-driven passive linear displacement. I. Basic properties. Journal of Neurophysiology, 77(6), 318077192.

Kaas, J. H., Gharbawie, O. A., & Stepniewska, I. (2013). Cortical networks for ethologically relevant behaviors in primates. American Journal of Primatology, 75(5), 407–414.

Kawano, K., Sasaki, M., & Yamashita, M. (1980). Vestibular input to visual tracking neurons in the posterior parietal association cortex of the monkey. Neuroscience Letters, 17(1–2), 55–60.

Kobayashi, Y., & Amaral, D. G. (2000). Macaque monkey retrosplenial cortex: I. Three-dimensional and cytoarchitectonic organization. Journal of Comparative Neurology, 426(3), 339–365.

Kobayashi, Y., & Amaral, D. G. (2003). Macaque monkey retrosplenial cortex: II. Cortical afferents. Journal of Comparative Neurology, 466(1), 48–79.

Kolb, B., Buhrmann, K., McDonald, R., & Sutherland, R. J. (1994). Dissociation of the medial prefrontal, posterior parietal, and posterior temporal cortex for spatial navigation and recognition memory in the rat. Cerebral Cortex, 4(6), 664–80.

Komatsu, H., & Wurtz, R. H. (1988). Relation of cortical areas MT and MST to pursuit eye movements. I. Localization and visual properties of neurons. Journal of Neurophysiology, 60(2), 580–603.

Kondo, H., Saleem, K. S., & Price, J. L. (2005). Differential connections of the perirhinal and parahippocampal cortex with the orbital and medial prefrontal networks in macaque monkeys. Journal of Comparative Neurology, 493(4), 479–509.

Kravitz, D. J., Saleem, K. S., Baker, C. I., & Mishkin, M. (2011). A new neural framework for visuospatial processing. Nature Reviews. Neuroscience, 12(4), 217–230.

Kropff, E., Carmichael, J. E., Moser, M.-B., & Moser, E. I. (2015). Speed cells in the medial entorhinal cortex. Nature, 523(7561), 419–424.

Lakshminarasimhan, K. J., Petsalis, M., Park, H., DeAngelis, G. C., Pitkow, X., & Angelaki, D. E. (2017). A Dynamic Bayesian Observer Model Reveals Origins of Bias in Visual Path Integration. bioRxiv 191817. https://doi.org/10.1101/191817

Liu, J., & Newsome, W. T. (2002). Functional Organization of Speed Tuned Neurons in Visual Area MT. Journal of Neurophysiology, 89(1), 246–256.

Liu, S., Gu, Y., DeAngelis, G. C., & Angelaki, D. E. (2012). Choice-related activity and correlated noise in subcortical vestibular neurons. Nature Neuroscience, 16(1), 89–97.

Lu, X., & Bilkey, D. K. (2010). The velocity-related firing property of hippocampal place cells is dependent on self-movement. Hippocampus, 20(5), 573–583. https://doi.org/10.1002/hipo.20666

Maaswinkel, H., & Whishaw, I. Q. (1999). Homing with locale, taxon, and dead reckoning strategies by foraging rats: sensory hierarchy in spatial navigation. Behavioural Brain Research, 99(2), 143–52. Retrieved from http://www.ncbi.nlm.nih.gov/pubmed/10512581

Maciokas, J. B., & Britten, K. H. (2010). Extrastriate area MST and parietal area VIP similarly represent forward headings. Journal of Neurophysiology, 104(1), 239–47.

Maguire, E. A. (1998). Knowing Where and Getting There: A Human Navigation Network. Science, 280(5365), 921807–924.

Maier, A., Aura, C. J., & Leopold, D. A. (2011). Infragranular Sources of Sustained Local Field Potential Responses in Macaque Primary Visual Cortex. Journal of Neuroscience, 31(6), 1971–1980.

McNaughton, B. L., Mizumori, S. J., Barnes, C. A., Leonard, B. J., Marquis, M., & Green, E. J. (1994). Cortical representation of motion during unrestrained spatial navigation in the rat. Cerebral Cortex (New York, N. Y. : 1991), 4(1), 27–39.

Merchant, H., Battaglia-Mayer, A., & Georgopoulos, A. P. (2001). Effects of optic flow in motor cortex and area 7a. Journal of Neurophysiology, 86(4), 1937–1954.

Merchant, H., Battaglia-Mayer, A., & Georgopoulos, A. P. (2004). Neural Responses during Interception of Real and Apparent Circularly Moving Stimuli in Motor Cortex and Area 7a. Cerebral Cortex, 14(3), 314–331.

Mitchell, J. F., Sundberg, K. A., & Reynolds, J. H. (2007). Differential Attention-Dependent Response Modulation across Cell Classes in Macaque Visual Area V4. Neuron, 55(1), 131–141.

Mitzdorf, U., & Singer, W. (1979). Excitatory synaptic ensemble properties in the visual cortex of the macaque monkey: A current source density analysis of electrically evoked potentials. Journal of Comparative Neurology, 187(1), 71–83.

Morris, R., Petrides, M., & Pandya, D. N. (1999). Architecture and connections of retrosplenial area 30 in the rhesus monkey (macaca mulatta). European Journal of Neuroscience, 11(7), 2506–2518.

Motter, B. C., & Mountcastle, V. B. (1981). The functional properties of the light-sensitive neurons of the posterior parietal cortex studied in waking monkeys: foveal sparing and opponent vector organization. Journal of Neuroscience, 1(1), 3–26.

Nandy, A. S., Nassi, J. J., & Reynolds, J. H. (2017). Laminar Organization of Attentional Modulation in Macaque Visual Area V4. Neuron, 93(1), 235–246.

Nitz, D. (2009). Parietal cortex, navigation, and the construction of arbitrary reference frames for spatial information. Neurobiology of Learning and Memory, 91(2), 179–185.

Nitz, D. A. (2006). Tracking Route Progression in the Posterior Parietal Cortex. Neuron, 49, 747–756.

Oleksiak, A., Klink, P. C., Postma, A., van der Ham, I. J. M., Lankheet, M. J., & van Wezel, R. J. a. (2011). Spatial summation in macaque parietal area 7a follows a winner-take-all rule. Journal of Neurophysiology, 105(3), 1150–8.

Pandya, D. N., & Seltzer, B. (1982). Intrinsic connections and architectonics of posterior parietal cortex in the rhesus monkey. Journal of Comparative Neurology, 204(2), 196–210.

Parron, C., & Save, E. (2004). Evidence for entorhinal and parietal cortices involvement in path integration in the rat. Experimental Brain Research, 159(3), 349–359.

Phinney, R. E., & Siegel, R. M. (2000). Speed selectivity for optic flow in area 7a of the behaving macaque. Cerebral Cortex (New York, N.Y. : 1991), 10(4), 413–421.

Pouget, A., & Sejnowski, T. J. (1997). Spatial Transformations in the Parietal Cortex Using Basis Functions. Journal of Cognitive Neuroscience, 9(2), 222–237.

Qi, X.-L. L., Katsuki, F., Meyer, T., Rawley, J. B., Zhou, X., Douglas, K. L., & Constantinidis, C. (2010). Comparison of neural activity related to working memory in primate dorsolateral prefrontal and posterior parietal cortex. Frontiers in Systems Neuroscience, 4, 12.

Quraishi, S., Heider, B., & Siegel, R. M. (2007). Attentional modulation of receptive field structure in area 7a of the behaving monkey. Cerebral Cortex, 17(8), 1841–1857.

Raffi, M., & Siegel, R. M. (2007). A functional architecture of optic flow in the inferior parietal lobule of the behaving monkey. PLoS ONE, 2(2), e200.

Rawley, J. B., & Constantinidis, C. (2010). Effects of task and coordinate frame of attention in area 7a of the primate posterior parietal cortex. Journal of Vision, 10(1), 12.1–16.

Read, H. L., & Siegel, R. M. (1997). Modulation of responses to optic flow in area 7a by retinotopic and oculomotor cues in monkey. Cerebral Cortex, 7(7), 647–661.

Robinson, D. L., Bowman, E. M., & Kertzman, C. (1995). Covert orienting of attention in macaques. II. Contributions of parietal cortex. Journal of Neurophysiology, 74(2), 698–712.

Robinson, S., & Bucci, D. J. (2012). Damage to posterior parietal cortex impairs two forms of relational learning. Frontiers in Integrative Neuroscience, 6, 45.

Rockland, K. S., & Van Hoesen, G. W.. (1999) Some temporal and parietal cortical connections converge in CA1 of the primate hippocampus. Cerebral Cortex, 9(3), 232–7.

Rosenbaum, R. S., Ziegler, M., Winocur, G., Grady, C. L., & Moscovitch, M. (2004). “I have often walked down this street before”: fMRI studies on the hippocampus and other structures during mental navigation of an old environment. Hippocampus, 14(7), 826–835.

Rozzi, S., Calzavara, R., Belmalih, A., Borra, E., Gregoriou, G. G., Matelli, M., & Luppino, G. (2006). Cortical connections of the inferior parietal cortical convexity of the macaque monkey. Cerebral Cortex, 16(10), 1389–1417.

Saleem, K. S., & Logothetis, N. K. (2012). A Combined MRI and Histology Atlas of the Rhesus Monkey Brain in Stereotaxic Coordinates. Academic Press. USA.

Sargolini, F., Fyhn, M., Hafting, T., McNaughton, B. L., Witter, M. P., Moser, M.-B., & Moser, E. I. (2006). Conjunctive Representation of Position, Direction, and Velocity in Entorhinal Cortex. Science, 312(2006), 758–763.

Sato, N., Sakata, H., Tanaka, Y. L., & Taira, M. (2006). Navigation-associated medial parietal neurons in monkeys. Proceedings of the National Academy of Sciences of the United States of America, 103(45), 17001–6.

Sato, N., Sakata, H., Tanaka, Y. L., & Taira, M. (2010). Context-dependent place-selective responses of the neurons in the medial parietal region of macaque monkeys. Cerebral Cortex, 20(4), 846–858.

Save, E., Guazzelli, A., & Poucet, B. (2001). Dissociation of the effects of bilateral lesions of the dorsal hippocampus and parietal cortex on path integration in the rat. Behavioral Neuroscience, 115(6), 1212–23.

Save, E., & Poucet, B. (2000). Involvement of the hippocampus and associative parietal cortex in the use of proximal and distal landmarks for navigation. Behavioural Brain Research, 109, 195–206.

Save, E., & Poucet, B. (2008). Role of the parietal cortex in long-term representation of spatial information in the rat. Neurobiology of Learning and Memory, 91, 172–178.

Schlack, A., Hoffmann, K. P., & Bremmer, F. (2002). Interaction of linear vestibular and visual stimulation in the macaque ventral intraparietal area (VIP). European Journal of Neuroscience, 16(10), 1877–1886.

Schroeder, C. (1998). A spatiotemporal profile of visual system activation revealed by current source density analysis in the awake macaque. Cerebral Cortex, 8(7), 575–592.

Self, M. W., van Kerkoerle, T., Supèr, H., & Roelfsema, P. R. (2013). Distinct Roles of the Cortical Layers of Area V1 in Figure-Ground Segregation. Current Biology, pp. 2121–2129.

Seltzer, B., Cola, M. G., Gutierrez, C., Massee, M., Weldon, C., & Cusick, C. G. (1996). Overlapping and nonoverlapping cortical projections to cortex of the superior temporal sulcus in the rhesus monkey: Double anterograde tracer studies. Journal of Comparative Neurology, 370(2), 173–190.

Seltzer, B., & Pandya, D. N. (1986). Posterior parietal projections to the intraparietal sulcus of the rhesus monkey. Experimental Brain Research, 62(3), 459–469.

Shelton, A. L., & Gabrieli, J. D. E. (2002). Neural correlates of encoding space from route and survey perspectives. The Journal of Neuroscience, 22(7), 2711–7.

Siegel, R. M., & Read, H. L. (1997). Analysis of optic flow in the monkey parietal area 7a. Cerebral Cortex, 7(4), 327–346.

Snyder, L. H., Grieve, K. L., Brotchie, P., & Andersen, R. A. (1998). Separate body- and world-referenced representations of visual space in parietal cortex. Nature, 394(6696), 887–891.

Song, Y. H., Kim, J. H., Jeong, H. W., Choi, I., Jeong, D., Kim, K., & Lee, S. H. (2017). A Neural Circuit for Auditory Dominance over Visual Perception. Neuron, 93(4), 940–954.

Spiers, H. J., & Maguire, E. A. (2006). Thoughts, behaviour, and brain dynamics during navigation in the real world. NeuroImage, 31(4), 1826–1840.

Spiers, H. J., & Maguire, E. A. (2007). Neural substrates of driving behaviour. NeuroImage, 36(1), 245–255.

Steinmetz, M. A., Motter, B. C., Duffy, C. J., & Mountcastle, V. B. (1987). Functional properties of parietal visual neurons: radial organization of directionalities within the visual field. Journal of Neuroscience, 7(1), 177–191.

Steinmetz, M. a, Connor, C. E., Constantinidis, C., & McLaughlin, J. R. (1994). Covert attention suppresses neuronal responses in area 7a of the posterior parietal cortex. Journal of Neurophysiology, 72(2), 1020–1023.

Takahashi, K., Gu, Y., May, P. J., Newlands, S. D., DeAngelis, G. C., & Angelaki, D. E. (2007). Multimodal coding of three-dimensional rotation and translation in area MSTd: comparison of visual and vestibular selectivity. Journal of Neuroscience, 27(36), 9742–56.

Traverse, J., & Latto, R. (1986). Impairments in route negotiation through a maze after dorsolateral frontal, inferior parietal or premotor lesions in cynomolgus monkeys. Behavioural Brain Research. 20(2):203–15.

Van Essen, D. C., Felleman, D. J., DeYoe, E. A., Olavarria, J., & Knierim, J. (1990). Modular and hierarchical organization of extrastriate visual cortex in the macaque monkey. Cold Spring Harbor Symposia on Quantitative Biology, 1990;55:679–96.

Ventre, J., & Faugier-Grimaud, S. (1989). Projections du cortex pariétal sur le complexe vestibulaire du macaque. Rev Neurol (Paris), 145(8–9), 646–651.

Vogt, B. A., & Pandya, D. N. (1987). Cingulate cortex of the rhesus monkey: II. Cortical afferents. J Comp Neurol, 262(2), 271–289.

Whitlock, J. R., Pfuhl, G., Dagslott, N., Moser, M. B., & Moser, E. I. (2012). Functional Split between Parietal and Entorhinal Cortices in the Rat. Neuron, 73(4), 789–802.

Wilber, A. A., Skelin, I., Wu, W., & McNaughton, B. L. (2017). Laminar Organization of Encoding and Memory Reactivation in the Parietal Cortex. Neuron, 95(6), 1406–1419.

Wolbers, T., Weiller, C., & Büchel, C. (2004). Neural foundations of emerging route knowledge in complex spatial environments. Cognitive Brain Research, 21(3), 401–411.

Yamawaki, N., Radulovic, J., & Shepherd, G. M. G. (2016). A Corticocortical Circuit Directly Links Retrosplenial Cortex to M2 in the Mouse. Journal of Neuroscience, 36(36), 9365–9374.

Yang, Y., Liu, S., Chowdhury, S. A., DeAngelis, G. C., & Angelaki, D. E. (2011). Binocular Disparity Tuning and Visual-Vestibular Congruency of Multisensory Neurons in Macaque Parietal Cortex. Journal of Neuroscience 31(49), 17905–17916.

Yoon, K., Buice, M. a, Barry, C., Hayman, R., Burgess, N., & Fiete, I. R. (2013). Specific evidence of low-dimensional continuous attractor dynamics in grid cells. Nature Neuroscience, 16(8), 1077–1084.

Yushkevich, P. A., Piven, J., Hazlett, H. C., Smith, R. G., Ho, S., Gee, J. C., & Gerig, G. (2006). User-guided 3D active contour segmentation of anatomical structures: Significantly improved efficiency and reliability. NeuroImage, 31(3), 1116–1128.

Zhang, T., & Britten, K. H. (2010). The responses of VIP neurons are sufficiently sensitive to support heading judgments. Journal of Neurophysiology, 103(4), 1865–1873.

Zipser, D., & Andersen, R. A. (1988). A back-propagation programmed network that simulates response properties of a subset of posterior parietal neurons. Nature, 331, 679–684.

